# *C. elegans* DSB-3 Reveals Conservation and Divergence among Protein Complexes Promoting Meiotic Double-Strand Breaks

**DOI:** 10.1101/2021.05.14.444243

**Authors:** Albert W. Hinman, Hsin-Yi Yeh, Baptiste Roelens, Kei Yamaya, Alexander Woglar, Henri-Marc G. Bourbon, Peter Chi, Anne M. Villeneuve

## Abstract

Meiotic recombination plays dual roles in the evolution and stable inheritance of genomes: recombination promotes genetic diversity by reassorting variants, and it establishes temporary connections between pairs of homologous chromosomes that ensure for their future segregation. Meiotic recombination is initiated by generation of double-strand DNA breaks (DSBs) by the conserved topoisomerase-like protein Spo11. Despite strong conservation of Spo11 across eukaryotic kingdoms, auxiliary complexes that interact with Spo11 complexes to promote DSB formation are poorly conserved. Here, we identify DSB-3 as a DSB-promoting protein in the nematode *Caenorhabditis elegans*. Mutants lacking DSB-3 are proficient for homolog pairing and synapsis but fail to form meiotic crossovers. Lack of crossovers in *dsb-3* mutants reflects a requirement for DSB-3 in meiotic DSB formation. DSB-3 concentrates in meiotic nuclei with timing similar to DSB-1 and DSB-2 (predicted homologs of yeast/mammalian Rec114/REC114), and DSB-1, DSB-2, and DSB-3 are interdependent for this localization. Bioinformatics analysis and interactions among the DSB proteins support the identity of DSB-3 as a homolog of MEI4 in conserved DSB-promoting complexes. This identification is reinforced by colocalization of pairwise combinations of DSB-1, DSB-2, and DSB-3 foci in structured illumination microscopy images of spread nuclei. However, unlike yeast Rec114, DSB-1 can interact directly with SPO-11, and in contrast to mouse REC114 and MEI4, DSB-1, DSB-2 and DSB-3 are not concentrated predominantly at meiotic chromosome axes. We speculate that variations in the meiotic program that have co-evolved with distinct reproductive strategies in diverse organisms may contribute to and/or enable diversification of essential components of the meiotic machinery.

**Significance Statement:** Faithful inheritance of chromosomes during meiosis depends on the formation and repair of double-strand DNA breaks (DSBs), which are generated through the activity of a topoisomerase-like protein known as Spo11. Spo11 exhibits strong conservation throughout eukaryotes, presumably reflecting constraints imposed by its biochemical activity, but auxiliary proteins that collaborate with Spo11 to promote and regulate DSB formation are less well conserved. Here we investigate a cohort of proteins comprising a complex required for meiotic DSB formation in *Caenorhabditis elegans*, providing evidence for both conservation with and divergence from homologous complexes in other organisms. This work highlights the evolutionary malleability of protein complexes that serve essential, yet auxiliary, roles in fundamental biological processes that are central to reproduction.

## Introduction

Meiotic recombination is important for two reasons. It promotes genetic diversity by reassorting traits, and it is important for creating temporary attachments between pairs of homologous chromosomes that are necessary for their future segregation at the meiosis I division. Recombination is initiated by the programmed introduction of DNA double-strand breaks (DSBs) (1). Some DSBs are repaired by a mechanism that leads to the formation of crossovers (COs) between homolog pairs, and the remaining DSBs are repaired as noncrossover products, thereby restoring genome integrity. Although DSBs are required for CO formation, they may lead to genomic instability if they are not repaired or are repaired erroneously. Thus, DSB formation in meiotic cells is governed by regulatory and surveillance mechanisms that function to ensure that enough DSBs are created to guarantee a CO on each homolog pair while limiting excess DSBs that may endanger the genome (2). Without appropriate DSB formation and repair, COs may fail to form between homologs during meiotic prophase, resulting in unattached homologs (univalents) that mis-segregate during the meiotic divisions, leading to aneuploidy in the resulting progeny.

Meiotic DSB formation is catalyzed by Spo11, a topoisomerase-like protein homologous to the catalytic A subunit of archaeal class VI topoisomerases that is well conserved across eukaryotic kingdoms (3–6). The mechanism of DNA breakage involves formation of a covalent linkage between Spo11 protein DNA, analogous to a key intermediate in the topisomerase reaction (1). Despite identification of structural and mechanistic conservation between Spo11 and TopVIA more than 20 years ago, however, counterparts of the archaeal TopVIB subunit that partner with Spo11 in “Spo11 core complexes” were not recognized until much later, reflecting substantial divergence both from TopVIB and among their eukaryotic orthologs (7–9).

DSB formation also depends on multiple additional factors that play critical roles in determining the location, timing, levels, and regulation of DSB formation (2). Several of these auxiliary DSB-promoting factors, including Rec114, Mei4 and Mer2, were originally discovered through genetic screens in *Saccharomyces cerevisiae* designed to identify genes required for initiation of recombination (10, 11). In contrast to the high level of conservation observed for Spo11, but similar to the other subunits of the Spo11 core complex, many auxiliary DSB protein such Rec114, Mei4 and Mer2 are poorly conserved at the primary sequence level (1). Indeed, high levels of sequence divergence had prevented identification of Rec114, Mei4 and Mer2 homologs outside of budding yeast until non-standard bioinformatics approaches were applied (12, 13). Homologs of Rec114 and Mei4 that are required for meiotic recombination have now been identified in several species, including *Mus musculus* (13–15), *Schizosaccharomyces pombe* (11, 16, 17), and *Arabidopsis thaliana* (18, 19). Proteins that were independently discovered based on roles in meiotic recombination in the ascomycete *Sordaria macrospora* (Asy1) and in the nematode *Caenorhabditis elegans* (DSB-1 and DSB-2) were also subsequently identified as putative Rec114 homologs (12, 20, 21), but Mei4 homologs were not yet identified in these organisms.

Several studies have established that DSB auxiliary factors Rec114 and Mei4 work closely together with each other and with Mer2 to promote meiotic DSB formation. Physical interactions among these proteins and their orthologs have been demonstrated for several organisms (13, 22–26), and coimmunoprecipitation experiments in *M.* musculus have further confirmed that these proteins interact with one another *in vivo* in a meiotic context (15). Recent biochemical analyses have shown that Rec114 and Mei4 together form individual complexes with a stoichiometry of 2 Rec114 molecules for every 1 Mei4 molecule and have further suggested that these complexes may self-assemble into large molecular condensates on chromatin during meiotic progression (27). In both *S. cerevisiae* and *M. musculus,* all three proteins have been reported to localize together in foci on meiotic prophase chromosomes (15, 22, 23, 26). Further, mouse REC114 and MEI4 and the Mer2 homolog IHO1 all localize predominantly at the meiotic chromosome axis (15, 26), contributing to the idea that they act as an intermediary between chromosome organization and DSB formation. Consistent with this view, chromatin immunoprecipitation experiments in both *S. cerevisiae* and *S. pombe* have shown that these proteins interact with both axis-enriched DNA sequences and with DSB sites (25, 28–30). Additionally, *S. cerevisiae* Rec114 and Mei4 have been found to interact with the Rec102 and Rec104 subunits that together comprise the TopVIB-like component of the Spo11 core complex (9, 23). Together these findings implicate Rec114-Mei4 in recruiting Spo11 to the meiotic chromosome axis.

*C. elegans* DSB-1 and DSB-2, while clearly implicated in meiotic DSB formation, were difficult to recognize as Rec114 homologs owing to high sequence divergence (12, 20, 21). Further, *C. elegans* differs from yeast and mice regarding the relationships between DSB formation and meiotic chromosome organization. Whereas DSB-dependent recombination intermediates are required to trigger assembly of the synaptonemal complex (SC) between homologous chromosomes in yeast and mice, *C. elegans* can achieve full synapsis between aligned homologs even in the absence of DSB formation (6). Thus, there are substantial differences in the cellular environments in which DSB-promoting complexes have evolved and function in different organisms.

In our current work, we identify DSB-3 as a protein that partners with DSB-1 and DSB-2 to promote SPO-11-dependent meiotic DSB formation in *C. elegans*. We demonstrate a requirement for DSB-3 in promoting the DSBs needed for CO formation, and we show that DSB-3 becomes concentrated in germ cell nuclei during the time when DSBs are formed, in a manner that is interdependent with DSB-1 and DSB-2. Through a combination of bioinformatics, interaction data, and colocalization analyses, we identify DSB-3 as a likely Mei4 homolog and establish DSB-1-DSB-2-DSB-3 as functional counterpart of the Rec114-Mei4 complex. Despite homology and a shared role in promoting DSB formation, we uncover surprising differences between the *C. elegans* DSB-1-DSB-2-DSB-3 and the REC114-MEI4 complexes observed in mice, notably that *C. elegans* DSB-1, DSB-2 and DSB-3 are distributed broadly on chromatin rather than becoming concentrated preferentially on chromosome axes. This work highlights the evolutionary malleability of protein complexes that serve essential, yet auxiliary, roles in meiotic recombination. Rapid diversification of such proteins may reflect a relaxation of constraints enabled by changes in another aspect of the reproductive program, or alternatively, they may reflect a capacity of alterations in such proteins to have an immediate impact on reproductive success.

## Results

### Identification of *dsb-3* as a gene required for the formation of meiotic crossovers

The *dsb-3(me6ts)* allele was isolated in a genetic screen for mutants exhibiting a <u>h</u>igh incidence of <u>m</u>ales among the progeny of self-fertilizing hermaphrodites, *i.e.,* the “Him” phenotype, which is indicative of errors in segregation of X chromosomes during meiosis. *me6ts* mutant hermaphrodites exhibit temperature-sensitive meiotic defects affecting both autosomes and X chromosomes (Table 1, Figure 1A). Whereas inviable embryos and males (XO) are rare among the self-progeny of wild-type hermaphrodites (XX), *dsb-3(me6ts)* mutant hermaphrodites raised at the non-permissive temperature of 25°C produce 28% inviable embryos, and 17% of their surviving progeny are males. Further, DAPI staining of chromosomes in oocytes at diakinesis, the last stage of meiotic prophase, revealed a defect in chiasma formation in the *dsb-3(me6ts)* mutant, reflecting an underlying defect in crossover formation (see below). Whereas wild-type oocyte nuclei consistently exhibit 6 pairs of homologous chromosomes connected by chiasmata (bivalents), oocyte nuclei in the *dsb-3(me6ts)* mutant exhibited a mixture of bivalents and unattached (achiasmate) chromosomes (univalents), with the incidence of univalents increasing with time.

**Figure 1.**
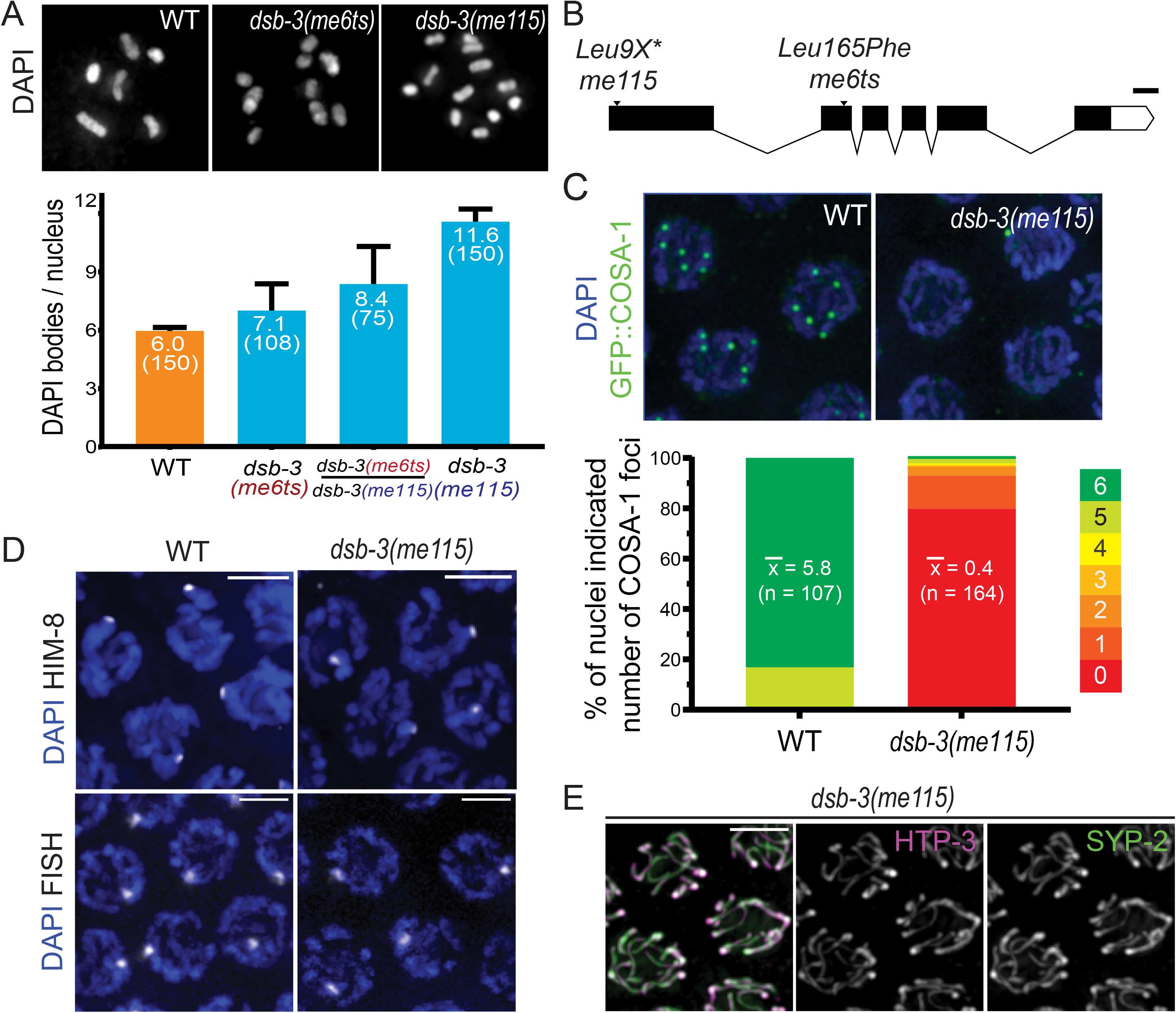
Identification of *dsb-3* as a gene required for the formation of meiotic crossovers. (A) Representative images of DAPI-stained diakinesis-stage oocyte nuclei from adult worms of the indicated genotypes fixed at 1 day post L4. Left: WT nucleus with six DAPI bodies corresponding to six pairs of homologs connected by chiasmata (bivalents). Middle: *dsb-3(me6ts)* nucleus with 9 DAPI bodies (3 bivalents and 6 univalents). Right: *dsb-3(me115)* nucleus with 12 DAPI bodies (all univalents). Below: Graphs show quantification of the mean number of DAPI bodies/ nucleus; error bars indicate standard deviation, and numbers in parentheses indicate the numbers of nuclei assayed. Assays for WT and for *dsb-3(me115)* homozygotes were performed at 20°C; assays for *dsb-3(me6ts)* homozygotes and *dsb-3(me6ts)/dsb-3(me115)* heterozygotes were performed at 25°C. (B) Schematic showing the *dsb-3* gene structure, with the positions and nature of mutations used in this work; white boxes represent UTR sequences, black boxes represent exons, lines indicate introns. Scale bar indicates 100 bp. (C) Top: Whole-mount immunofluorescence images of GFP::COSA-1 foci, which correspond to the single CO site on each homolog pair, in nuclei at the late pachytene stage. WT nuclei have 6 GFP::COSA-1 foci, while foci are reduced or absent entirely in the *dsb-3(me115)* mutant. Below: Stacked bar graphs showing the distribution of GFP::COSA-1 foci counts in nuclei from WT and *dsb-3(me115)* mutants. Mean numbers of GFP::COSA-1 foci per nucleus are indicated, with the numbers of nuclei assayed in parentheses. (D) Homolog pairing assayed by immunofluorescence of X-Chromosome pairing center binding protein HIM-8 (Top) or fluorescence *in situ* hybridization FISH detecting a 1Mbp segment of chromosome II (Bottom) in pachytene nuclei of whole-mount gonads. A single focus is observed in each nucleus, indicating successful pairing between the homologs. Scale bar is 3.2 µm. (E) Immunofluorescence image of SC components in late pachytene nuclei in a whole-mount gonad from the *dsb-3(me115)* mutant. Axis protein HTP-3 and SC central region protein SYP-2 colocalize in continuous stretches between chromosome pairs, indicating successful synapsis. Scale bar is 3.2 µm.

**Table 1.** Quantitation of embryo viability, male frequency and diakinesis karyotypes. Strains used in and/or created for this study were evaluated for the indicated parameters. Numbers of DAPI bodies were evaluated for worms fixed and stained at 24 hours post L4 stage. Analysis of AV913 and corresponding controls were conducted at two temperatures based on the temperature-sensitive nature of the *dsb-3(me6ts)* mutant.

| Strain | Genotype | Incubated Temperature | Average Brood Size ± SD<br>(number of broods) | Percent Dead Eggs | Percent Males | Average DAPI Bodies/Nucleus ± SD<br>(number of oocytes) | |
| --- | --- | --- | --- | --- | --- | --- | --- |
|  |  |  |  |  |  | 24 h post L4 | 48 h post L4 |
| N2 | WT | 20° | 264 ± 27 (7) | 0.4 | 0 | 6.0 ± 0.2 (92) | 5.9 ± 0.3 (102) |
| N2 | WT | 25° | 291 ± 22 (7) | 2 | 0 | 5.9 ± 0.3 (110) | 5.9 ± 0.4 (99) |
| AV913 | *dsb-3(me6ts)* IV | 20° | 274 ± 27 (7) | 0.1 | 0 | 6.0 ± 0.2 (144) | 5.9 ± 0.3 (111) |
| AV913 | *dsb-3(me6ts)* IV | 25° | 198 ± 47 (7) | 28.2 | 17.3 | 7.0 ± 1.4 (108) | 9.9 ± 1.3 (99) |
| AV1095 | *dsb-3(me115)* IV | 20° | 156 ± 77 (16) | 99.4 | 24.9 | 11.6 ± 0.6 (150) | |
| AV1029 | *meSi7[sun1p::dsb-3::gfp::sun1 3'UTR unc-119+] II; dsb-3(me115)* IV | 20° | 150 ± 23 (8) | 5 | 3.7 | 6.1 ± 0.4 (207) | |
| AV1102 | *dsb-3(me125)[3xflag::dsb-3] dsb-1(me124)[3xha::dsb-1]* IV | 20° | 243 ± 34 (7) | 2.1 | 2.5 | 6.0 ± 0.2 (213) | |
| AV1115 | *dsb-2(me132)[3xha::dsb-2]* II | 20° | 193 ± 61 (8) | 2.2 | 0.5 | 6.0 ± 0.2 (150) | |

Mapping and sequencing identified a missense mutation at genomic position IV: 7758710 (WS279) as the likely causal mutation responsible for the *dsb-3(me6ts)* mutant phenotype (see Materials and Methods); this mutation results in a Leu165Phe substitution in the previously uncharacterized protein C46A5.5. CRISPR/Cas9 genome editing was used to introduce multiple stop codons early into the first exon of *C46A5.5*, thereby creating the null allele *me115* (Figure 1B). *me115* fails to complement *dsb-3(me6ts)* (Figure 1A), confirming the identity of *C46A5.5* as *dsb-3*.

Analysis of the *dsb-3(me115)* null mutant indicates that the DSB-3 protein is required for the formation of meiotic crossovers between all six pairs of homologous chromosomes. *dsb-3(me115)* mutant hermaphrodites produced 99% inviable embryos, and 25% of their surviving progeny were male, reflecting mis-segregation of autosomes and X chromosomes (Table 1). Diakinesis oocytes of *dsb-3(me115)* mutant hermaphrodites exhibited an average of 11.6 ± 0.6 DAPI-stained bodies, indicating of a lack of chiasmata connecting all six homolog pairs (Figure 1A). A severe defect in crossover formation in the *dsb-3(me115)* mutant was also revealed using GFP::COSA-1 as a cytological marker of crossover-designated sites in late pachytene nuclei ((31); Figure 1C). Whereas 6 GFP::COSA-1 foci (1 per homolog pair) were consistently observed in late pachytene nuclei of control worms, GFP::COSA-1 foci were absent from most late pachytene nuclei in the *dsb-3(me115)* mutant.

Since pairing and assembly of the synaptonemal complex between homologous chromosomes are prerequisites for the formation of crossovers during *C. elegans* meiosis, we evaluated whether these features were impaired in the *dsb-3(me115)* mutant. We assessed pairing using fluorescence *in-situ* hybridization (FISH) for a 1 Mbp segment of Chromosome II and immunostaining for HIM-8, a C2H2 zinc-finger DNA-binding protein that concentrates on the Chromosome X pairing center (32, 33), demonstrating that *dsb-3(me115)* mutants are proficient for homolog paring (Figure 1D). Further, immunostaining for the axial element protein HTP-3 and the synaptonemal complex central region protein SYP-2 (34, 35) revealed fully synapsed chromosomes in early pachytene nuclei in the *dsb-3(me115)* mutant, indicating successful SC assembly (Figure 1E). Together these data indicate that the DSB-3 protein is dispensable for pairing and synapsis, and point to a role for this protein in the DNA events of recombination.

During *C. elegans* meiosis, failure to form crossover recombination intermediates between one or more chromosome pairs prolongs the early pachytene stage of meiotic prophase, reflecting the operation of a “crossover assurance” checkpoint (20, 21, 36). Consistent with the observed deficit of interhomolog crossovers, *dsb-3(me115)* mutant gonads display an extended zone of nuclei exhibiting phosphorylation of nuclear envelope protein SUN-1, a marker of crossover assurance checkpoint activation (Supplemental Figure 1A).

### DSB-3 is required for meiotic double-strand break formation

Meiotic recombination is initiated through the formation of DSBs by the conserved topoisomerase-like protein SPO-11 (3, 6). Following formation, these DSBs are then processed to enable the loading of the DNA strand-exchange protein RAD-51 (37, 38). RAD-51 foci thus mark the sites of recombination intermediates that can be assayed as a proxy for successful initiation of meiotic recombination (35, 39). We observed a strong decrease in the number of RAD-51 foci in *dsb-3(me115)* mutants relative to wild type (Figure 2A), suggesting that fewer DSBs are being created in these mutants or that there is a failure to load RAD-51 at DSB sites.

**Figure 2.**
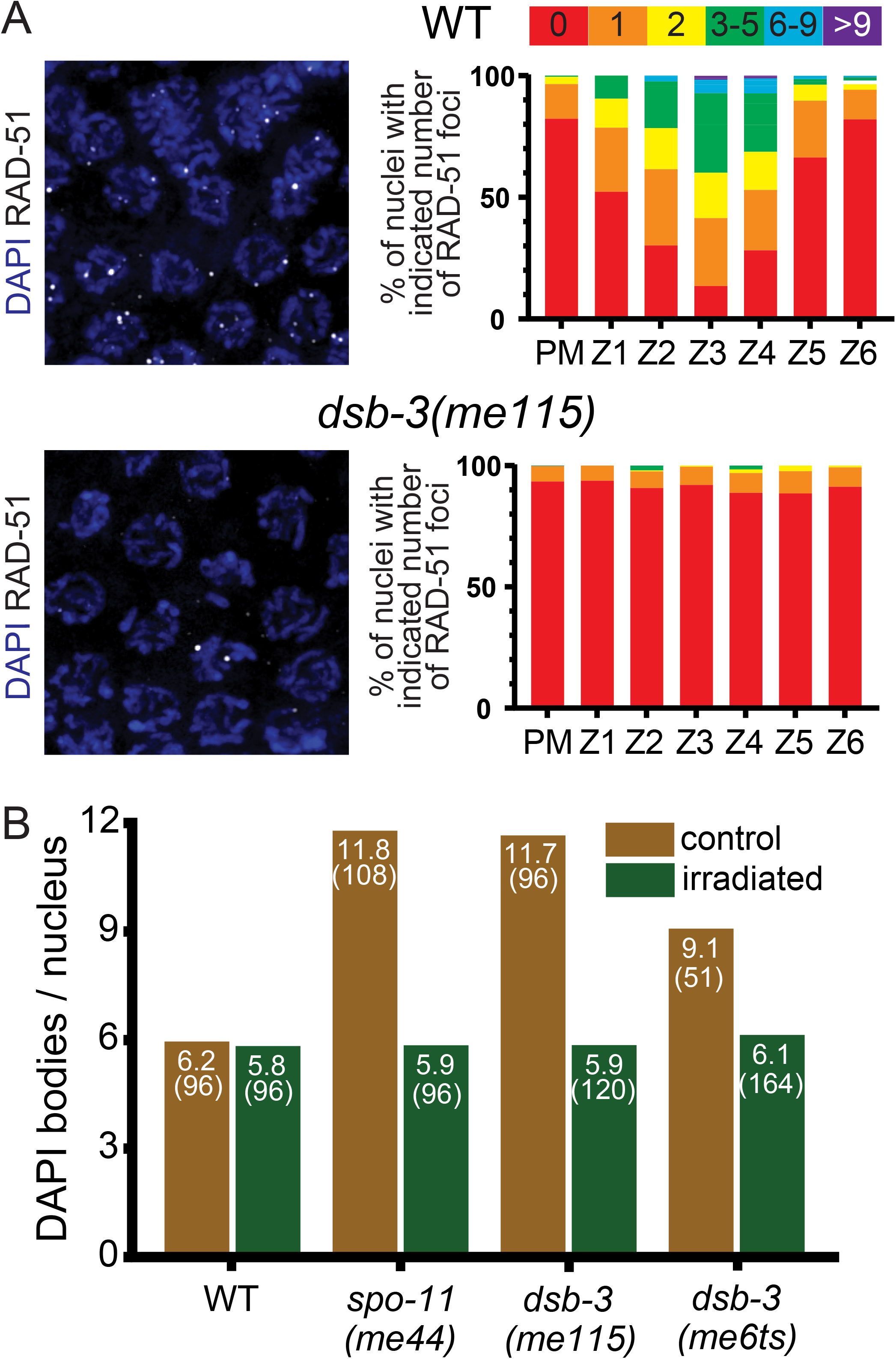
DSB-3 is required for meiotic double-strand break formation. (A) Left: Immunofluorescence images of RAD-51 foci in early pachytene nuclei in whole-mount gonads. RAD-51 foci mark sites of processed DSBs and are strongly reduced in the *dsb-3(me115)* mutant. Right: Quantification of RAD-51 foci in whole-mount gonads (at least three gonads were scored per genotype). Gonads were divided into seven zones: the premeiotic zone (PM), which includes all nuclei prior to the transition zone (where nuclei enter meiotic prophase), and six consecutive equal-sized zones encompassing the region of the gonad from the transition zone to the end of the pachytene stage. (B) Rescue of chiasma formation in *dsb-3* mutants by γ-irradiation induced DNA breaks. Graph showing the average numbers of DAPI bodies present in diakinesis-stage oocytes of worms exposed to 5 kRad of γ-irradiation at 20 hours post L4, and un-irradiated age-matched controls, fixed and stained with DAPI 18-20 hours post irradiation.

To determine whether fewer endogenous DSBs was the defect responsible for the observed reduction in RAD-51 foci, we used γ-irradiation to introduce ectopic DSBs to test whether such breaks are sufficient to restore crossover formation. Similar approaches were taken with other DSB-defective mutants in *C. elegans,* such as *dsb-1*, *dsb-2*, and *spo-11* (6, 20, 21). Young adult *dsb-3(me115)* and *dsb-3(me6ts)* hermaphrodites (alongside wild-type and *spo-11(me44)* controls) were exposed to 5000 rad of γ-irradiation and subsequently assayed for crossover formation through DAPI staining of chromosomes in oocyte nuclei. We observed a full rescue of normal DAPI-body counts after irradiation (Figure 2B), suggesting that the *dsb-3* mutants are specifically defective in DSB formation.

### DSB-3 is concentrated in DSB-competent nuclei and is interdependent with DSB-1 and DSB-2

Consistent with its role in promoting meiotic DSB formation, we find that DSB-3 localizes to germ cell nuclei during the time when meiotic DSBs are formed. To assess DSB-3 localization in the germ line, we generated a transgenic strain that expresses a DSB-3::GFP fusion protein in the germ line in the *dsb-3(me15)* null mutant background (*meSi7[sun-1p::dsb-3::gfp::sun13’UTR]* II; *dsb-3(me115)* IV); based on assessment of progeny viability and DAPI bodies in diakinesis oocytes, we infer that this DSB-3::GFP fusion protein is largely functional in promoting meiotic recombination (Table 1). Immunolocalization experiments in whole-mount dissected gonads show that DSB-3::GFP becomes concentrated in germ cell nuclei within the transition zone, soon after entry into meiotic prophase (Figure 3A). The DSB-3::GFP immunofluorescence signal is strongest in early pachytene nuclei, then declines sharply in mid-pachytene, albeit with a few outlier nuclei in the late pachytene region of the gonad retaining a strong DSB-3 signal. This pattern of appearance and disappearance of DSB-3 from germ cell nuclei is similar to the patterns observed for the double-strand break promoting proteins DSB-1 and DSB-2 ((20, 21) and Figure 3A), and corresponds to the timing when nuclei are competent for meiotic DSB formation.

**Figure 3.**
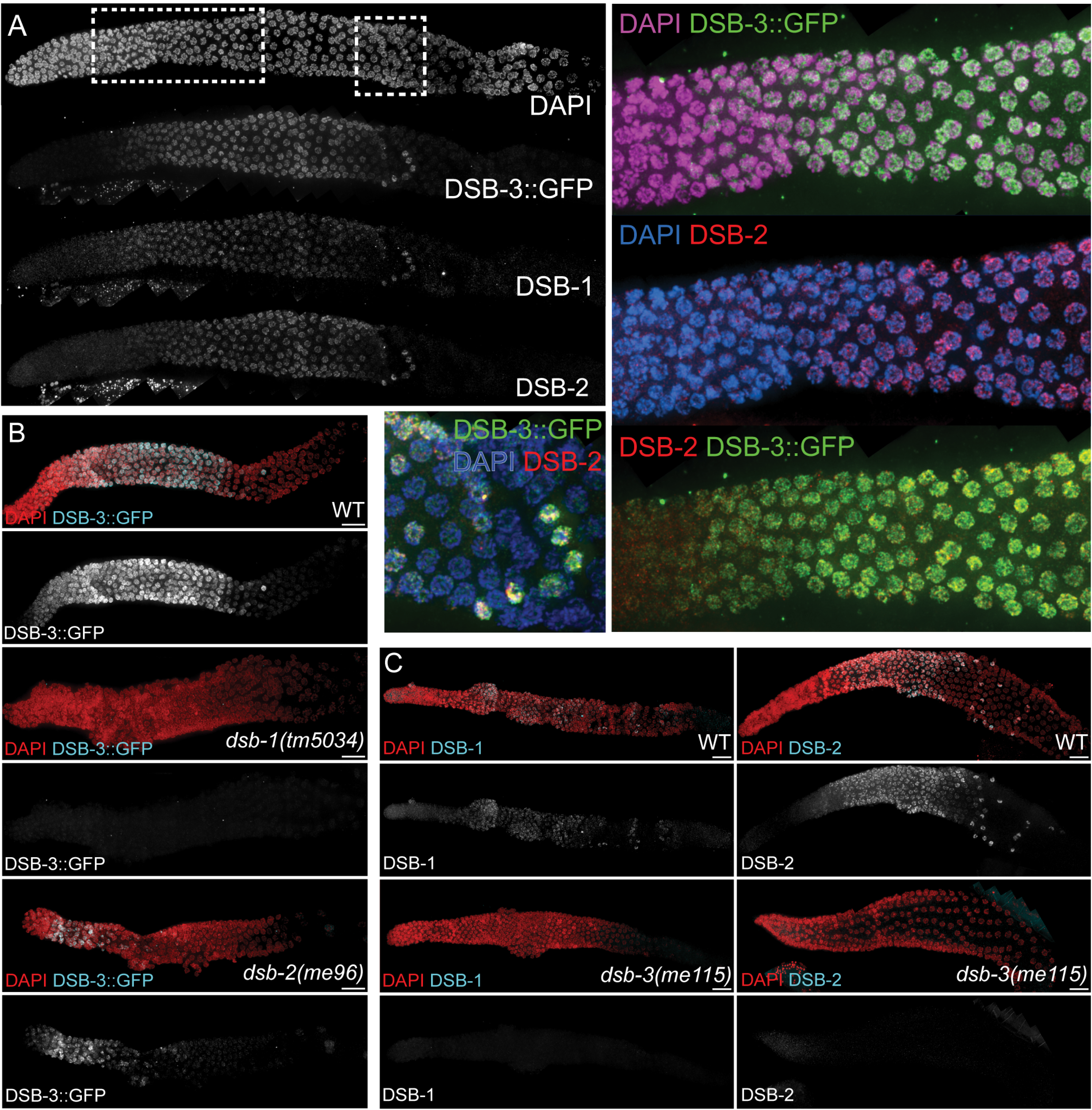
DSB-3 is concentrated in DSB-competent nuclei and is interdependent with DSB-1 and DSB-2. (A) Immunofluorescence image of a whole-mount hermaphrodite gonad (from distal tip to end of pachytene) stained with DAPI and antibodies detecting DSB-3::GFP, DSB-1, and DSB-2. Meiotic progression proceeds from left to right, as in the rest of the images for this figure. DSB-3::GFP becomes concentrated in germ cell nuclei within the transition zone (rectangular inset - top, soon after entry into meiotic prophase. The DSB-3 immunofluorescence signal is strongest in early pachytene nuclei, then declines sharply in mid-pachytene, albeit with a few outlier nuclei (square inset) in the late pachytene region of the gonad retaining a strong DSB-3 signal. This pattern of appearance and disappearance of DSB-3 from germ cell nuclei is similar to the patterns observed for the double-strand break promoting proteins DSB-1 and DSB-2. (B) Immunofluorescence images of a whole-mount hermaphrodite gonads stained with DAPI and antibody detecting DSB-3::GFP. DSB-3::GFP signal is not detected in the *dsb-1(null)* mutant, and is strongly reduced and limited to a smaller region of the gonad in the *dsb-2(null)* mutant. (C) Immunofluorescence images of gonads stained for DSB-1 (left) or DSB-2(right), showing that DSB-1 and DSB-2 immunofluorescence signals are not detected in the *dsb-3(me115)* mutant. Scale bars = 16.2 µm.

The DSB-3, DSB-2, and DSB-1 proteins are not only abundant in the same nuclei during early meiotic prophase, but they are also interdependent for this immunolocalization. Previous studies had demonstrated interdependence for DSB-1 and DSB-2 (20, 21), with loss of the DSB-2 immunofluorescence signal in a *dsb-1* null mutant (which lacks meiotic DSB-promoting activity) and reduction of DSB-1 immunofluorescence signal in a *dsb-2* null mutant (which retains a low residual level of DSB-promoting activity). Similarly, we found that DSB-3::GFP immunofluorescence signal was abolished in *dsb-1* null mutant germ lines (Figure 3B). Likewise, DSB-3::GFP immunofluorescence signal was reduced in *dsb-2* null mutant germ lines and was restricted to a few rows of nuclei in the transition zone and very early pachytene regions of the gonad (Figure 3B). Conversely, DSB-1 and DSB-2 immunostaining were lost in *dsb-3(me115)* mutant germ lines (Figure 3C). Collectively, these data indicate that DSB-3, DSB-2, and DSB-1 are interdependent for proper localization to germ cell nuclei, indicating that they function together in promoting meiotic DSB formation.

Together, our data demonstrating 1) a similar requirement in promoting DSB formation, 2) concentration in the same nuclei, and 3) interdependence for localization and/or abundance in meiotic nuclei are all consistent with DSB-3 functioning in a protein complex together with DSB-1 and DSB-2 to promote the formation of SPO-11-dependent meiotic DSBs.

### Evidence that DSB-1, DSB-2 and DSB-3 form a complex homologous to the yeast and mammalian Rec114-Mei4 complexes

Although the initial PSI-BLAST searches conducted for DSB-1 and DSB-2 had not identified homologs outside of *Caenorhabditis* (20, 21), DSB-1 and DSB-2 were subsequently identified as likely distant homologs of the Rec114 meiotic DSB-promoting proteins from fungi and mammals (12). This identification was enabled using an approach involving PSI-BLAST searches initiated using sequence alignments, in combination with scanning for patterns of similarity in predicted secondary structure, to identify short signature motifs (SSMs) in poorly conserved proteins (40). We obtained additional support for the assignment of DSB-1 and DSB-2 as Rec114 homologs using the Phyre2 structure prediction server (41), which identified medium confidence (#6 hit, 40.3%) and low confidence (#19 hit, 11.9%) alignments between the N-terminal domains of DSB-2 and DSB-1 and the N-terminal domain (where the identified SSMs are located) of the solved structure of mouse Rec114 (42). We therefore used an alignment driven PSI-BLAST approach similar to that described above to identify DSB-3 as a putative homolog of Mei4 (see Materials and Methods, Figure 4A, Supplemental Figure 2), which is required for meiotic DSB formation in yeast and mice and forms a complex with Rec114 (13–15, 22, 24, 43). Based on local amino acid composition and relative position in the protein sequence, three of the 6 SSMs previously identified in Mei4 homologs from diverse species (SSMs #1, 4 and 6) are well supported in the *Caenorhabditis* DSB-3 orthologs, while the other three SSM are less well conserved.

**Figure 4.**
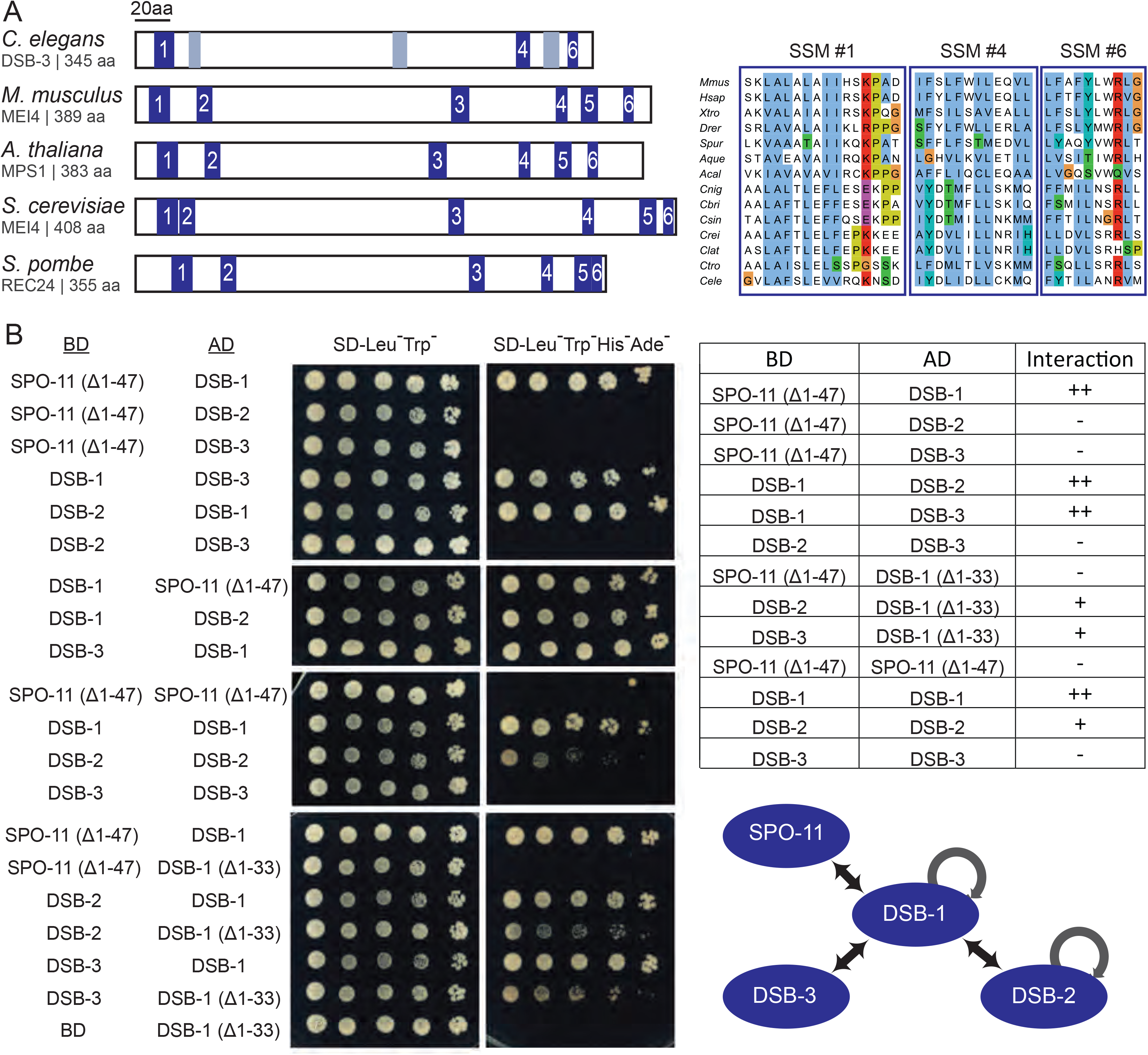
Evidence that DSB-1, DSB-2 and DSB-3 form a complex homologous to the yeast and mammalian REC114-MEI4 complexes. (A) Left: Schematic diagram depicting the positions of six short signature motifs (SSMs, blue boxes) previously defined for Mei4 homologs in diverse species, indicating the three SSMs (1, 4 and 6) that are most strongly supported in *C. elegans* DSB-3; gray boxes indicate positions that potentially correspond to SSMs 2, 3 and 5 based on a multiple sequence alignment (MSA) with vertebrate and marine invertebrate homologs of similar size, but are less well conserved. Right: Aligned sequences of SSMs 1, 4 and 6 from *Mus musculus, Homo sapiens, Xenopus tropicalis, Danio rerio*, *Strongylocentrotus purpuratus, Amphimedon queenslandica, Aplysia californica* and the following nematodes of the genus *Caenorhabditis*: *C. nigoni, C. briggsae, C. sinica, C. remanei, C. latens, C. tropicalis, C. elegans*. SSMs are cropped from an MSA generated in JalView (80) using the ClustalX coloring scheme. For additional information, see Supplemental Figure 2 and Materials and Methods. (B) Yeast two-hybrid assay revealing protein-protein interactions among the DSB-1, −2, and −3 proteins and between DSB-1 and SPO-11. Potential interactions between proteins fused with the GAL4 activation domain (AD) and proteins fused with the GAL4 DNA binding domain (BD) were assayed by growth on media lacking histidine and adenine. A construct producing an N-terminal truncation of SPO-11 lacking the first 47 amino acids (Δ1-47) was used for these analyses, as severe auto-activation was observed for full-length SPO-11. Negative controls showing lack of auto-activation for the constructs used are presented in Supplemental Figure 2. In addition to experiments using constructs expressing full-length DSB-1, some experiments used a construct expressing an N-terminally truncated DSB-1 lacking the first 33 amino acids (Δ1-33). A schematic summarizing the identified interactions is shown on the bottom right.

To complement these *in silico* analyses, we used yeast two-hybrid (Y2H) assays to establish a network of interactions among the DSB-1, DSB-2, and DSB-3 proteins and SPO-11, the protein that catalyzes DSB formation (Figure 4B, Supplemental Figure 2). Y2H interactions were detected between DSB-1 and DSB-2 and between DSB-1 and DSB-3, consistent with an ability of these proteins to form complexes. Homotypic interactions were also detected both for DSB-1 and for DSB-2. In addition to the interactions detected among the putative Rec114 and Mei4 homologs, DSB-1 also interacted with SPO-11 in the Y2H assay.

We note that a truncated version of DSB-1 lacking the N-terminal 33 amino acids loses the ability to interact with SPO-11 but retains its ability to associate with DSB-2 and DSB-3. This suggests that the interactions between DSB-1 and SPO-11 and the interactions between DSB-1 and DSB-2 or DSB-3 may be mediated, at least in part, by different parts of the DSB-1 protein.

### DSB-3, DSB-2, and DSB-1 colocalize in meiotic nuclei

To complement our genetic, bioinformatic, and Y2H evidence that DSB-3, DSB-2, and DSB-1 function together as components of conserved protein complexes to promote DSB formation, we investigated their colocalization using Structured Illumination Microscopy (SIM) on spread preparations of meiotic nuclei (Figure 5). For most of these analyses, we used a moderate nuclear spreading protocol (44), coupled with SIM imaging to provide improved spatial resolution below the limits of standard light microscopy (45). This approach enabled detection of these proteins as chromosome-associated foci. To facilitate co-staining of protein pairs for these colocalization analyses, we used CRISPR/Cas9 gene editing to create strains expressing HA or FLAG tagged versions of the DSB proteins from the endogenous loci (Table 1), and we detected the proteins using indirect immunofluorescence.

**Figure 5.**
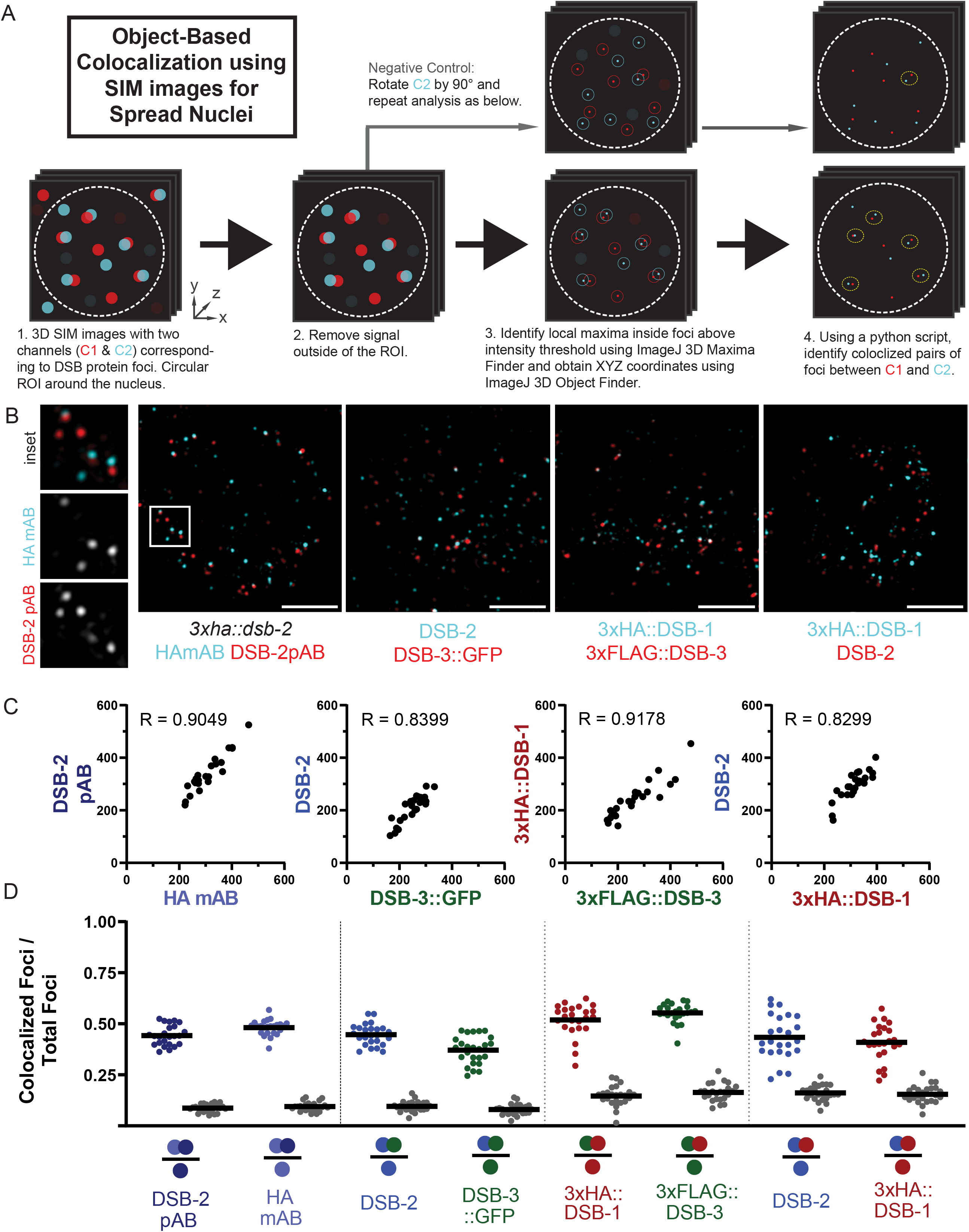
DSB-3, DSB-2, and DSB-1 colocalize in meiotic nuclei. (A) Schematic summarizing the object-based image analysis pipeline used to assess colocalization of foci in SIM immunofluorescence images of partially spread nuclei (see Materials and Methods and Supplemental Figure 3 for more details). (B) Representative SIM immunofluorescence images of individual spread nuclei stained with antibodies targeting the indicated DSB proteins; each panel depicts a single Z slice from a 3D image stack. The following experiments are represented in order from left to right: mouse monoclonal HA and rabbit polyclonal DSB-2 antibodies, both detecting the same 3xHA::DSB-2 tagged protein in a *3xha::dsb-2* II strain; chicken GFP and rabbit DSB-2 antibodies, detecting DSB-3::GFP and DSB-2 in a *meSi7[dsb-3::gfp]* II; *dsb-3(me115)* IV strain; mouse HA and rabbit FLAG antibodies, detecting 3xHA::DSB-1 and 3xFLAG::DSB-3 in a *3xflag::dsb-3 3xha::dsb-1* IV strain; and mouse HA and rabbit DSB-2 antibodies, detecting 3xHA::DSB-1 and DSB-2 in a *3xflag::dsb-3 3xha::dsb-1* IV strain. Scale bar is 2 µm. (C) Quantification of DSB protein foci for the indicated pairwise combinations; each data point represents the numbers of foci for the two analyzed channels in a single nucleus. Spearman R values reported indicate that numbers of the two types of foci are strongly correlated within nuclei. (D) Graphs showing the fraction of foci of a given type (indicated by a single colored circle in the denominator in each schematic below the horizontal axis) that are colocalized with the other type of focus analyzed in that same experiment (colocalizing foci are represented by two colored circles in the numerator). For each pair of focus types analyzed, two sets of experimental analyses (represented by colored data points) and paired negative controls (represented by grey data points) are presented.

We used the image analysis pipeline outlined in Figure 5A (described in more detail in Supplemental Figure 3) to identify DSB protein foci and to conduct object-based colocalization analyses to assess the degree of colocalization detected for pairwise combinations of the imaged DSB proteins within individual nuclei. As negative controls, we generated virtual nuclei in which the second channel in each combination was rotated by 90° in XY, resulting in virtual composite images in which DSB protein foci are modified in location, but numbers, sizes, and intensity distributions of foci remain unaltered. Collectively, our analyses indicate that the DSB-3, DSB-2, and DSB-1 proteins strongly colocalize with each other in meiotic prophase nuclei.

Analysis of all three pairwise combinations of DSB-1, DSB-2, and DSB-3 foci are presented in Figure 5B-D and Supplemental Figure 4. For all three pairs, numbers of foci for the two channels detected in each nucleus were strongly correlated, consistent with expectations for components of the same protein complex. Further, substantial colocalization was observed for each pair. For example, we found that 45 ± 5% of DSB-2 foci colocalized with DSB-3::GFP foci, and conversely, that 37 ± 7% of DSB-3::GFP foci colocalized with DSB-2 foci. In contrast, negative control coincidental colocalization values were 10 ± 3% and 8 ± 3%, respectively.

Similarly, 52 ± 8% of 3xFLAG::DSB-3 foci colocalized with 3xHA::DSB-1 foci, and conversely 55% ± 5% of 3xHA::DSB-1 foci colocalized with 3xFLAG::DSB-3 foci. Likewise, 43 ± 11% of 3xHA::DSB-1 foci colocalized with DSB-2 foci, and conversely 41 ± 9% of DSB-2 foci colocalized with 3xHA::DSB-1 foci.

Although substantial colocalization was observed for all pairwise combination of DSB-1, DSB-2 and DSB-3 foci, the fraction of colocalization may seem lower than might be anticipated for proteins comprising the same protein complex. We note, however, that incomplete colocalization has been similarly observed for the Rec114 and Mei4 proteins in both budding yeast and mouse meiocytes (22, 26). One possible explanation is that only a subset of these protein molecules occur together in complexes, while other molecules exist separately within the nucleus; however, this explanation is not easily reconciled with the observed interdependence among these proteins. Another possibility is that the observed degree of colocalization reflects limitations on our ability to detect all of the DSB-1, DSB-2 and DSB-3 target molecules that are present. *e.g.* because of isoforms that lack epitopes or because the complexes and/or their components may be organized in a manner that makes some epitopes inaccessible to detection reagents.

This latter possibility is supported by data from an experiment in which we assessed colocalization for fluorescent foci representing separate epitopes on the same protein, 3xHA::DSB-2, expressed from the endogenous *dsb-2* locus. Specifically, we used a mouse monoclonal antibody (mAB) against the HA epitope and rabbit polyclonal (pAB) antibodies raised against the C-terminal 100 amino acids of the DSB-2 protein. The numbers of foci for the two channels detected in each nucleus were strongly correlated (Figure 5C), as expected for foci representing the same target molecule. However, colocalization was again incomplete, in both directions: 44 ± 5% of DSB-2 pAB foci colocalized with HA mAB foci, and conversely, 48 ± 2% of HA mAB foci colocalized with DSB-2 pAB foci. This incomplete colocalization of HA mAB and DSB-2 pAB fluorescent signals supports the conclusion that a subset of epitopes on DSB-2 proteins present in the nucleus were not detected in these experiments.

For the DSB-2 - DSB-3::GFP combination, we also conducted a colocalization analysis on “super-spread” nuclei, in which chromosomes were dispersed over an area 6-10 times larger than that of an unperturbed nucleus (46) (Materials and Methods and Supplemental Figure S5). The numbers of foci detected using this approach were 3-5 times higher than the numbers observed in our analysis of partial spreads, but a similar degree of colocalization was detected: 55 ± 16% of DSB-2 foci colocalized with DSB-3::GFP foci, and conversely, 44 ± 10% of DSB-3::GFP foci colocalized with DSB-2 foci. The observation of larger numbers of foci with a comparable degree of colocalization suggests the possibility that groups of DSB protein complexes may be split into smaller cohorts by the super-spread procedure.

### The Presence and Colocalization of DSB-3, DSB-2, and DSB-1 Is Not Confined to the Meiotic Chromosomal Axis

Previous chromatin immunoprecipitation experiments in *S. cerevisiae* have shown that Mei4 and Rec114 are enriched at DNA sequences that are also enriched for meiosis-specific axis proteins Hop1 and Red1 (28, 30). Moreover, *M. musculus* Mei4 and Rec114 were found to colocalize cytologically on the axes of meiotic prophase chromosomes (14, 15, 26). Based on these observations, it has been proposed that the Rec114-Mei4 complex primarily functions at the chromosome axes. Strikingly, DSB-1, DSB-2, and DSB-3 foci are not enriched at the chromosome axis during *C. elegans* meiosis. Rather, we find that most foci are detected away from the axis, in the associated chromatin loops (Figure 6A). To quantify axis association, we first segmented images by creating axis masks for each nucleus that corresponded to the pixels containing the immunofluorescence signal derived from the axis protein HTP-3 (Supplemental Figure 3B), then for each DSB protein tested, we identified the subset of foci, termed “axis-associated foci” for which some or all of the pixels coincided with the axis mask. This approach indicated that only 15-30% of DSB protein foci detected in our analyses overlapped with chromosome axis signal, indicating that on spread chromosomes from meiotic nuclei. the preponderance of DSB protein foci detected were not associated with the meiotic chromosome axes.

**Figure 6.**
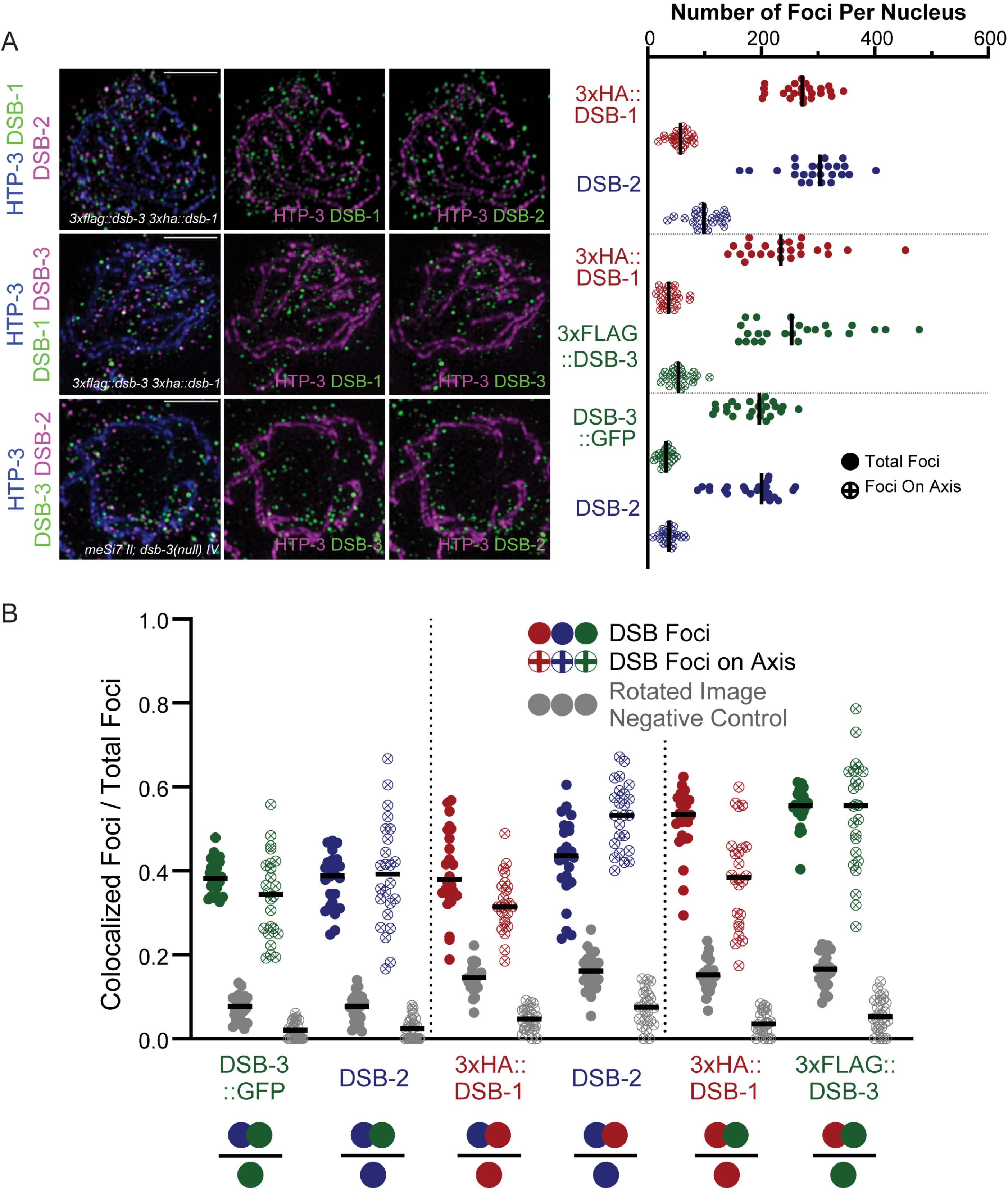
The Presence and Colocalization of DSB-3, DSB-2, and DSB-1 are Not Confined to the Meiotic Chromosomal Axis. (A) Left: representative maximum-intensity projections of SIM images, representing the middle third of Z-stacks collected for the depicted spread nuclei. Nuclei were stained with antibodies targeting DSB proteins and the axis component HTP-3. Scale bar is 2 µm. Right: graph showing quantification of numbers of DSB protein foci that colocalize with the axis signal, as well as the total numbers of DSB protein foci identified in the analyzed nuclei. The data indicate that the majority of foci do not colocalize with the axis. (B) Graphs depicting the fractions of Colocalized Foci / Total Foci for DSB protein foci of the indicated types (represented as in Figure 5), reported for axis-associated foci and for total nuclear foci analyzed within the same data sets. The filled dots represent colocalization fractions for all DSB protein foci of a given type within the nucleus, while dots with crosses represent the colocalization fractions for the subset of DSB protein foci of the indicated type that colocalize with the axis.

We also assessed whether DSB protein foci associated with the chromosome axis might exhibit a higher degree of colocalization with their DSB protein partners relative to the level of colocalization observed for the full set of foci within the nucleus (Figure 6B). However, this analysis did not reveal any consistent enrichment of colocalization for DSB protein foci that were linked to the chromosome axis. Thus, while components of the meiotic chromosome axis do have roles in promoting and regulating SPO-11 dependent DSB forming activity in *C. elegans* meiosis(47–49), these roles do not appear to be mediated by concentrating DSB-1-DSB-2-DSB-3 complexes in close proximity to the chromosome axis.

## Discussion

Initiation of meiotic recombination by programmed DSB formation is an ancient and conserved feature of the meiotic program that predates divergence of plants, animals and fungi. Thus, it is not surprising that Spo11, the protein directly responsible for catalyzing DSB formation, is strongly conserved across kingdoms, given constraints imposed by its requirement to interact with and perform chemistry on DNA. However, many additional proteins required for DSB formation had been identified in the yeast system, but plant and metazoan homologs of these auxiliary DSB proteins had long eluded detection by standard BLAST analyses. The barrier to detection of homologs outside fungi was eventually breached using secondary structure prediction coupled to MAFFT alignment and phylogenomically-oriented PSI-BLAST searches (13), which identified characteristic SSMs for putative Mei4 and Rec114 homologs; moreover, the veracity of these predictions was borne out by demonstration of meiotic roles in mouse mutants (13–15).

Auxiliary proteins involved in DSB formation during *C. elegans* meiosis were identified independently based on analysis of meiotic mutants ((20, 21), this work). However, recognition of these proteins as distant homologs of conserved DSB-promoting factors came later, after their functional importance in DSB formation was already established (12). Identification of DSB-1 and DSB-2 as Rec114 homologs was further solidified by alignments of the predicted structure of DSB-1 and DSB-2 with the solved structure of mouse REC114 ((42); this work). Likewise, our initial identification of *C. elegans* DSB-3 as a factor important for meiotic DSB formation was similarly based on functional data. The identity of DSB-3 as a putative Mei4/MEI4 ortholog was derived computationally from alignments and collinearity of SSMs among metazoan homologs, and this identification was reinforced by demonstration of Y2H interactions, colocalization and interdependence with Rec114 homologs DSB-1 and DSB-2. Thus, despite a high degree of divergence at the amino acid sequence level, our data collectively support the conclusion that DSB-1, DSB-2 and DSB-3 together form complexes that are the functional counterpart of Rec114-Mei4 complexes.

Having established conservation among the auxiliary complexes that promote the DSB-forming activity of Spo11, our analyses also reveal interesting differences. First, whereas yeast and mice each have only a single Rec114/REC114 ortholog, nematodes in the Caenorhabditis genus each have 2 paralogs, indicating duplication and divergence in the parental lineage. DSB-1 and DSB-2 are neither identical to nor functionally interchangeable with each other, as DSB formation is strongly reduced in *dsb-2* mutants and eliminated in *dsb-1* mutants. Interestingly, recent biochemical analyses indicate a 2 Rec114 : 1 Mei4 stoichiometry of the yeast complex (27). The interdependence of DSB-1 and DSB-2 for nuclear enrichment, in combination with the colocalization observed in chromosome spreads (this work) suggest that the *C. elegans* complexes may typically contain one DSB-1 subunit and one DSB-2 subunit (rather than two identical subunits). However, there is low residual DSB-promoting activity present in *dsb-2* null mutants, suggesting that complexes with two DSB-1 subunits may form and be partially functional when DSB-2 is absent. While the data do support functional diversification of the *C. elegans* Rec114 paralogs, however, how this diversification came about and/or how and why it persisted remain unknown.

A second apparent distinction between *C. elegans* DSB-1, DSB-2 and DSB-3 and their mouse counterparts is the observed relationship to meiotic chromosome axes. Mouse REC114 and MEI4 are reported to localize predominantly at chromosome axes in spread preparations of meiotic prophase chromosomes, in a manner mediated by Mer2 homolog IHO1 (15, 26). This association with chromosome axes led to the proposal that a major role for the Rec114 – Mei4 complex is to recruit the Spo11 core complex to the axis to activate its DSB-promoting activity specifically in close proximity to the axis, where DSB repair predominantly occurs. In *C. elegans*, meiosis specific HORMAD proteins HTP-1 and HTP-3, which are major building blocks of the chromosome axis, are implicated in playing important roles in promoting and regulating meiotic DSB formation (47–49). HTP-3 is strictly required for DSB formation, and HTP-1 is required for normal levels of DSB activity. However, despite this requirement for axis components in DSB formation, our analysis here indicates that DSB-1-DSB-2-DSB-3 complexes are not preferentially enriched adjacent to axes. This may reflect lack of an apparent *Caenorhabditis* ortholog of IHO1, which is recruited to mouse meiotic chromosomes through a direct interaction with HORMAD1.

Given the observed lack of enrichment of DSB-1, DSB-2 and DSB-3 at the axes, the role(s) of *C. elegans* meiotic HORMAD proteins in promoting DSB formation may not be strictly limited to recruiting the homologs of Rec114 and Mei4 to chromosomes. One possibility is that the *C. elegans* HORMADs might be involved in activating the subset of DSB-1-DSB-2-DSB-3 complexes that do occur in close proximity to the axis. Alternatively, the role(s) of the *C. elegans* axis proteins might be indirect, *e.g.* assembly of the axis might potentially signal successful formation of chromosome structure that is proficient for meiotic DSB repair, thereby licensing the nucleus that it is safe to proceed with DSB formation. The possibility of this signaling scenario is strengthened by prior work demonstrating a role for *C. elegans* HORMAD proteins in a signaling process that sustains activity of protein kinase CHK-2, a master regulator of multiple processes during meiosis, including nuclear enrichment of DSB-1 and DSB-2 (20, 21, 36, 50).

It is possible that the observed difference between mice and *C. elegans* regarding axis enrichment of Rec114-Mei4 complexes may be related to differences in spatial organization of recombination events in the genome and/or in coupling between DSB repair and homolog recognition. Meiotic recombination in mice occurs predominantly within 1-2 kb “hotspot’ regions, separated by larger (50-100kb) cold regions where the probability of recombination is very low. A similar “local hotspot” distribution of recombination events was not observed in *C. elegans* (for the portion of the genome studied) (51, 52), suggesting that different constraints are operating to dictate where DSBs may form. Further, there is substantial variation among organisms regarding their relative dependence on different mechanisms that promote pairwise alignment and synapsis between homologous chromosomes. In mouse meiosis, formation of early SPO11-dependent DSB repair intermediates appears to be the main mechanism of homology verification, required to trigger SC assembly and constraining it to occur strictly between aligned homologous chromosomes (53, 54). In contrast, in *C. elegans*, local *cis-*acting chromosomal domains known as pairing centers play a primary role in homolog recognition, and these are capable of promoting largely successful pairwise synapsis between homologs even in the absence of recombination (55). We speculate that differences in the constraints governing the genomic locations of DSBs and/or differences in dependence on DSBs for homology verification may have either contributed to, or been enabled by, diversification of meiotic DSB auxiliary protein complexes. We note that high divergence among essential components of key biological processes is a hallmark not only of the meiotic program, but of reproduction more generally (1, 56, 57). This likely reflects multiple underlying factors, including the potential for changes in such proteins to have an immediate impact on processes that directly affect fitness by modulating reproductive success.

A key question regarding the role(s) of the Rec114-Mei4 (or DSB-1-DSB-2-DSB-3) complexes is how exactly they are functioning to promote Spo11 activity. It was recently proposed that Rec114-Mei4 complexes function by forming large DNA-dependent biomolecular condensates that promote DSB activity by causing a high local concentration of Spo11 core complexes at presumptive DSB sites held adjacent to the chromosome axis (27). This model was proposed based on: 1) a large segment of the Rec114 protein exhibiting a high probability of disorder, 2) the ability of purified Rec114-Mei4 complexes to promote formation of DNA-dependent condensates *in vitro*, and 3) an ability of Rec114-Mei4 complexes to interact with and recruit the Spo11 core complex. While evaluating potential for *in vitro* condensation is outside the scope of the current study, we note that some of the above attributes may be shared with the *C. elegans* DSB-1-DSB-2-DSB-3 complex. First, based on the Y2H data, the DSB-1-DSB-2-DSB-3 complex is expected to be able to interact with SPO-11 complexes. However, in yeast, interactions occur between Rec114 and the Rec102 and Rec104 components (which together correspond to Top6BL) of the Spo11 core complex, whereas in *C. elegans,* a Top6BL homolog has not yet been identified, and DSB-1 can interact directly with SPO-11 itself in the Y2H assay. Second, the DSB-1 and DSB-2 proteins have long segments with predicted protein disorder scores that hover around 0.5 and include short segments scoring >0.5, leaving it ambiguous whether these might represent *bona fide* disordered regions. Third, the observation that higher numbers of DSB-2 and DSB-3 foci are detected in super-spread nuclei than in moderately-spread nuclei raises the possibility that these proteins might normally occur in larger groups within intact nuclei, potentially analogous to the condensates proposed to occur during yeast meiosis. In either system, future investigations aiming to test predictions of the condensation model will need to address the challenge of visualizing complex dynamic behavior *in vivo*.

## Materials and Methods

### C. elegans strains

Strains were cultured at 20℃ using standard nematode growth conditions (58) unless otherwise noted. Strains used in this study:

AV28 *dsb-3(me6ts)* IV

AV776 *spo-11(me44) IV / nT1[qIs51]* (IV;V)

AV818 *meIs8[gfp::cosa-1]* II; *cosa-1(tm3298)* III

AV913 *dsb-3(me6ts)* IV

AV958 *dsb-3(me6ts) dpy-20(e1282)* IV

AV994 *dpy-3(e184) dsb-3(me6ts)* IV

AV995 *dsb-3(me115) IV / nT1 (*IV;V)

* AV1029 *meSi7* [*sun1p::dsb-3::gfp::sun-1 3’UTR] II; dsb-3(me115) IV*

AV1045 *meSi7 II; dsb-3(me115) dsb-1(we11) / nT1 IV*

AV1081 *meSi7 dsb-2(me96) / mnC1 II; dsb-3(me115) IV*

AV1095 *dsb-3(me115) / tmC5 [F36H1.3(tmIs1220)]* IV

^ AV1102 *dsb-1(me124[3xha::dsb-1]) dsb-3(me125[3xflag::dsb-3]) IV*

^ AV1115 *dsb-2(me132)[3xha::dsb-2] II*

AV1132 *meIs8[gfp::cosa-1]* II*; cosa-1(tm3298)* III*; dsb-3(me115)* IV */ nT1* (IV;V)

Bristol N2 Wild type

* The transgene allowing expression of the DSB-3::GFP fusion protein was obtained using the Mos Single Copy Insertion strategy (MosSCI, (59)) using the *ttTi5605* insertion on chromosome II as a landing site. The donor plasmid, pBR253, was obtained by assembling fragments carrying the upstream promoter region of the *sun-1* gene, the *sun-1* downstream 3’UTR region, and the genomic sequence of *dsb-3* (coding exons and introns), together with a DNA fragment containing a version of GFP optimized for germline expression (60), into pBR49, a derivative of pCFJ350 modified to enable type IIs restriction/ligation cloning (61). The genomic fragments were obtained by PCR amplification of wild-type genomic DNA using the following primer pairs. The primers for the *sun-1* promoter were oBR840 (cgtcgatgcacaatccGGTCTCaCCTGatttccagatttcatcgtcggtttt) and oBR841 (agtggaatgtcagGGTCTCaCATaccgagtagatctggaagtttag). The primers for *dsb-3* CDS were: oBR836 (cgtcgatgcacaatccGGTCTCaTATGATCGAAATTACCGATGATGAGG) and oBR837 (agtggaatgtcagGGTCTCaCTCCATTGCTATATCTCTGTTGATTATCTAAAAAC) The primers for the *sun-1* 3’UTR were oBR842 (cgtcgatgcacaatccGGTCTCaTAAAaaacgccgtattattgttcctgc) and oBR843 (agtggaatgtcagGGTCTCaGTCAttagtaagttaaagctaaagttagcag). The GFP fragment was obtained by PCR amplification of pCFJ1848 (60), using oBR406 (cgatgcacaatccGGTCTCaGGAGGTGGATCATCCTCCACATCATCCT) and oBR407 (agtggaatgtcagGGTCTCaTTTATGGGGAAGTACCGGATGACG). Correct assembly of all fragments within the donor plasmids was verified by sequencing.

^ In order to perform pairwise colocalization experiments between DSB-1, DSB-2 and DSB-3, we created strains expressing endogenously-tagged versions of these proteins so each pair could be detected using compatible primary antibodies generated in different host organisms. For these strains, we used direct injection of Cas9 protein (PNAbio) complexed with single-guide RNA (sgRNA) (Dharmacon) using the protocol of (62). CRISPR targeting (crRNA) sequences were designed using Benchling (https://benchling.com/). Small single-stranded oligonucleotides (< 200 bp) were purchased (Integrated DNA Technologies) and used as the repair templates to generate the various tags and nonsense alleles. N2 worms (P0) were injected with the mix together with sgRNA and repair template for the *dpy-10* co-CRISPR marker (63). Rol F1s (carrying *dpy-10(Rol)* marker) were singled out, and a subset of F2 progeny was fixed and stained with DAPI (see below) to assess the phenotype of diakinesis nuclei for null alleles. From plates containing worms exhibiting univalents at diakinesis, the new mutations were recovered from siblings of the imaged worms and balanced by *nT1* IV or *tmc5* IV. Tagged alleles were confirmed by immunofluorescence staining (see below). All edits were confirmed by Sanger sequencing of PCR fragments amplified using primers designed to detect the edit event. The crRNAs used, description of the edits, and PCR sequencing primers used are included in Supplemental Table 1.

### Isolation, mapping, and genomic identification of the *dsb-3(me6ts)* mutation

*dsb-3(me6ts)* was isolated in a genetic screen for meiotic mutants exhibiting a high incidence of males as described in (64). After backcrossing (four times) to generate the AV913 strain, homozygous *me6ts* worms were subjected to whole-genome sequencing. DNA was extracted from ∼8 60mm confluent plates of N2 and AV913 gravid adult worms; worms were rinsed twice in M9 and resuspended in 10 mM EDTA and 0.1 M NaCl. Worms were then: pelleted; flash frozen in liquid nitrogen; resuspended in 450 μL of lysis buffer containing 0.1 M Tris, pH 8.5, 0.1 M NaCl, 50 mM EDTA, and 1% SDS plus 40 μL of 10 mg/mL proteinase K in TE (10 mM Tris, 1 mM EDTA), pH 7.4; vortexed; and incubated at 62°C for 45 min. Two successive phenol-chloroform extractions were performed using the Phase Lock gel tubes from Invitrogen, and DNA was precipitated with 1 mL of 100% ethanol plus 40 μL of saturated NH_4_Ac (5 M) and 1 μL of 20 mg/mL GlycoBlue. The DNA pellet was washed with 70% ethanol, air dried, and resuspended in 50 μL of TE, pH 7.4. Paired end libraries were prepared using the Nextera technology (Illumina), and sequencing was performed on an MiSeq sequencer (2 × 75 bp) through the Stanford Functional Genomics Facility. To analyze the genomic data, we used an analysis pipeline adapted from GATK’s recommended best practices (65–67). Reads were mapped to *C. elegans* reference genome (WBcel 235.82) using the Bowtie 2 software (68). Variant calling was performed using Haplotype Caller software from GATK, and lists from AV913 and N2 were compared to eliminate non-causal variants. The predicted effects of variants specific to AV913 were then annotated using SnpEff (69).

Initial genetic mapping experiments had placed *dsb-3(me6ts)* within 2 cM of *unc-5*, located at 1.78 cM on chromosome IV; the above sequence analysis identified several candidate mutations within this region. Additional mapping crosses located *dsb-3(me6ts)* to the left of *dpy-20* (at 5.22 cM) and near or to the left of *unc-24* (at 3.51 cM). Further, we found that *eDf18* (which deletes the region between 3.7-4.19 cM) complements *dsb-3(me6ts)*. Together, these experiments identified a G -> A transition at genomic position IV: 7758710 (WS279), in the second coding exon of the uncharacterized gene *C46A5.5*, as the likely causal mutation responsible for the *dsb-3(me6ts*) mutant phenotype.

### DAPI staining of oocyte chromosomes and Irradiation Assay

Numbers of DNA bodies present in diakinesis oocytes were assessed in intact adult hermaphrodites of the indicated ages, raised at the indicated temperatures, fixed in ethanol and stained with 4′,6-diamidino-2-phenylindole (DAPI) as in (70). This method underestimates the frequency of achiasmate chromosomes, as some univalents lie too close to each other to be resolved unambiguously.

To test for rescue of bivalent formation by exogenously derived DSBs, worms were exposed to 5,000 rad (50 Gy) of γ-irradiation using a Cs-137 source at 20 h post-L4 stage. Worms were fixed and stained at 18–20 h post-irradiation, and numbers of DAPI bodies were counted in oocyte nuclei in the −1 to −3 positions.

### Bioinformatic identification of homology between DSB-3 and Mei4

PSI Blast searches using the MPI BLAST server (71), initiated using an alignment of DSB-3 homologs from diverse roundworm species as the query, identified a putative *Brugia malayi* DSB-3 homolog. A subsequent round of PSI-BLAST searches, initiated using an alignment with the putative *B. malayi* homolog as the header sequence and initially focusing on the N-terminal portion of the protein, led to retrieval of plant and animal Mei4 homologs. Similarity in protein lengths and patterns of predicted secondary structure were prioritized over E-value considerations in selection of proteins chosen for the multiple sequence alignment presented in Supplemental Figure 2, which was generated using MAFFT Version 7.0 with gap opening penalty parameter set to 2.0 and offset value parament set to 0.125.

### Yeast Two-Hybrid Analysis

Full-length DSB-1, DSB-2, DSB-3, and N-terminally truncated SPO-11 (SPO-11Δ1-47), DSB-1 (DSB-1Δ1-33) ORFs were individually cloned into the *Bam*HI and *Pst*I sites of pBridge, and the *Bam*HI and *Xho*I sites of pGADT7 (Clontech) to generate fusion proteins with the N-terminal Gal4 DNA-binding domain (Gal4BD) or activation domain (Gal4AD). The PJ69-4A yeast strain (*MATa trp1-901 leu2-3,112 ura3-52 his3-200 gal4Δ gal8Δ GAL2-ADE2 LYS2::GAL1-HIS3 met2::GAL7-lacZ*) was co-transformed with the indicated pairs of constructs encoding Gal4BD and Gal4AD fusion proteins (and/or empty vector negative controls). Transformed cells expressing Gal4BD and Gal4AD fusion proteins were selected in SD-Leu^-^Trp^-^, a drop-out medium without leucine and tryptophan. Protein interactions were assayed by growing transformed cells for 5 days at 30°C on selective media lacking leucine, tryptophan, histidine, and adenine (SD-Leu^-^Trp^-^His^-^Ade^-^). Three independent repeats of each transformation were performed for all pairwise combinations. The full-length SPO-11 ORF was excluded from analysis of combinations as it exhibited autoactivation in negative control experiments.

### Immunofluorescence Methods

The following primary antibodies were used: mouse anti-HA (1:1000, Covance 16B12 clone), rabbit anti-FLAG (1:5000, Sigma Aldrich), rabbit anti-DSB-2 (1:5.000, (20)), guinea pig anti-DSB-1 (1:500, (21)), rabbit anti-GFP (1:200, (31)), guinea pig anti-HIM-8 (1:500, (32)), chicken anti-HTP-3 (1:400, (33)), rabbit anti-SYP-2 (1:200, (35)), rat anti-RAD-51 (1:500, (20)), guinea pig anti-SUN-1 S24pi (1:700, (72)), chicken anti-GFP (1:500, (A01694, Genscript)). Secondary antibodies were Alexa Fluor 488, 555 and 647-conjugated goat antibodies directed against the appropriate species (1:400, Life Technologies).

For immunofluorescence experiments involving whole mount gonads, dissection of gonads, fixation, immuno-staining and DAPI counterstaining were performed as in (48).

For experiments involving nuclear spreads, spreading was performed as in (73). The gonads of 20–100 adult worms were dissected in 10 µL Dissection solution (75% v/v Hank’s Balanced Salt Solution [HBSS, Life Technology, 24020-117] with 0.1% v/v Tween-20) on an ethanol-washed plain slide. 50 μL of spreading solution (32 μL of Fixative [4% w/v Paraformaldehyde and 3.2%–3.6% w/v Sucrose in water], 16 μL of Lipsol solution [1% v/v in water], 2 μL of Sarcosyl solution [1% w/v of Sarcosyl in water]) were added, and gonads were immediately distributed over the whole slide using a pipette tip. Slides were then left to dry at room temperature overnight, washed for 20 minutes in methanol at −20°C and rehydrated by washing 3 times for 5 minutes in PBS-T. A 20-minute blocking in 1% w/v BSA in PBS-T at room temperature was followed by overnight incubation with primary antibodies at room temperature (antibodies diluted in: 1% w/v BSA in PBS-T). Slides were washed 3 times for 5 minutes in PBS-T before secondary antibody incubation for 2 hours at room temperature. After PBS-T washes, the samples were mounted in Vectashield (Vector).

To dissect large quantities of *C. elegans* gonads for spreads, we employed an alternative method for disrupting worms, using a 125V ∼ 60Hz drill capable of achieving 1,600 rotations per minute. Briefly, we synchronized worms by using the standard bleaching protocol (74) and allowed worms grow to adulthood (L4 + 24 hours). The worms were then washed with dissection solution into a 1.7 mL Eppendorf tube suspended in an ethanol ice bath. The worms were then disrupted using a 1/64 inch bit on the drill with its maximum power by angling the drill bit against the Eppendorf tube wall. 3 μL aliquots were taken from the tube every 20 seconds and monitored microscopically until most of the gonads had been extruded from the worms during the drill-induced disruption.

### FISH experiments

Barcoded Oligopaint probes targeting a 1 Mb segment of chromosome II (genomic coordinates 11,500,001-12,500,001) were generated as in (46). Gonads from animals at 24 hours after L4 were dissected on a coverslip and fixed in 1% paraformaldehyde for 5 min. A slide (Superfrost Plus) was then placed on the coverslip and immersed in liquid N_2_. The sample was then incubated in −20°C methanol for 2 minutes and rehydrated by placing in PBST for at least 10 minutes. Next, the sample was incubated in 0.1 M HCl for 5 minutes and washed in PBST 3 times for 5 minutes each. The samples were then incubated for 5 minutes each in 2x SSCT (2x saline sodium citrate with 0.1% Tween) solutions with increasing concentrations of formamide: 0%, 5%, 10%, 25%, and 50%. The sample was then incubated in a prewarmed 42°C solution of 50% formamide in 2x SSCT for 1 hour. 2 μL of Oligopaint probe (1,000 ng/μL in dH2O) was diluted into 30 μL of hybridization solution (50% formamide, 10% Dextran Sulfate, 2x SSC, 0.1% Tween-20) for each slide. After 1 hour incubation, slides were taken out of the 50% formamide solution, wiped, and incubated in 95% ethanol for 5 minutes. Then, the probe hybridization solution was applied to the sample with a coverslip, and the sample was denatured for 10 minutes at 77°C on a heat block. After denaturing, the sample was incubated with the probe hybridization solution at 42°C overnight. The next day, samples were washed 2 times in 42°C 50% formamide in 2x SSCT for 30 minutes each, and the coverslip was removed from the slide. Then, the sample was incubated for 5 minutes each in solutions with decreasing concentrations of formamide in 2x SSCT: 25%, 10%, and 5%. Samples were then washed 2 times for 10 minutes each in 2x SSCT. The Oligopaint probes were visualized by hybridizing Cy3-labelled oligos (agctgatcgtggcgttgatg) to the Oligopaint probe barcode sequence. To do this, the Cy3-labelled probes (diluted 1:1000 in 25% ethylene carbonate in 2x SSC) were applied to the sample with a coverslip and incubated for 15 minutes. Then, the sample was washed and the coverslip was removed by incubating in 30% formamide solution in 2x SSCT for 3 min. The samples were then washed twice in 2x SSC and mounted in Vectashield.

For quantification of pairing between FISH signals, gonads were divided into 6 zones. Zone 1 corresponds to the distal tip region of the gonad with only premeiotic nuclei. The gonad region extending from the transition zone to the end of the pachytene stage was split up into 5 equally sized regions, Zones 2-6. The stitched image of the gonad was cropped into zones, peaks of FISH signals were identified using ImageJ plugin 3D Maxima Finder (75). Each identified peak was manually assigned to a nucleus, and distances between homologous signal peaks in the same nucleus were calculated.

### Image Acquisition

For spread nuclei, imaging, deconvolution, stitching and 3D-SIM reconstruction were performed as in (73). Spreading results in squashing of *C. elegans* germline nuclei from 5 to 1-2 µm in thickness. 3D-SIM images were obtained as 125 nm spaced Z-stacks, using a 100x NA 1.40 objective on a DeltaVison OMX Blaze microscopy system, 3D-reconstructed and corrected for registration using SoftWoRx. For display, images were projected using maximum intensity projection in ImageJ or SoftWoRx.

For imaging of whole-mount gonads. wide field (WF) images were obtained as 200 nm spaced Z-stacks, using a 100x NA 1.40 objective on a DeltaVison OMX Blaze microscopy system, deconvolved and corrected for registration using SoftWoRx. Subsequently, gonads were assembled using the “Grid/Collection” plugin (76) in ImageJ. For display, assembled gonads were projected using maximum intensity projection in ImageJ.

For display, contrast and brightness were adjusted in individual color channels using ImageJ.

### Quantification of RAD-51 Foci and COSA-1 Foci

For quantification of RAD-51 foci in whole-mount gonads, at least three gonads were counted per genotype. Gonads were divided into seven zones: the premeiotic zone (PM), which includes all nuclei prior to the transition zone (where nuclei enter meiotic prophase), and six consecutive equal-sized zones encompassing the region of the gonad from the transition zone to the end of the pachytene stage. For the GFP::COSA-1 experiments, foci were counted in nuclei within the last six cell rows of the gonad.

### Identification of DSB Protein Foci and Object-Based Colocalization Analysis

For Figures 5 and 6, images were analyzed using an object-based colocalization analysis pipeline that combined standard functions available in ImageJ in conjunction with a custom Python script. A detailed description of the colocalization analysis pipeline is presented in Supplemental Figure 3. For these analyses, 32-bit Z-stacks of SIM images of immunofluorescence signals for at least two different antibodies detecting DSB proteins (C1 and C2). were imported into ImageJ (77, 78) with the Fiji distribution (79). The signal maxima for each channel, identified as foci by the image analysis pipeline were qualitatively compared to the original image to verify accurate identification of foci.

For colocalization analysis of DSB-2 and DSB-3::GFP foci on super-spread nuclei (Supplemental Figure S5), the same pipeline was used, except that foci were analyzed within 3.43 x 3.43 μm square ROIs located entirely within the spread (1-3 ROIs per nucleus).

### Data and reagent availability

The original 32-bit individual nucleus ImageJ files, the segmented axis channel files, the identified peaks, the values used for and the output position files from the 3D Maxima Finder for each nucleus, the custom python script used to identify colocalization, and the resulting spreadsheet files showing colocalization data for each nucleus will be made available from the BioStudies Database, Accession Number S-BSST568.

Strains and primary images used in this research are available on request from A.M.V.

## Acknowledgments

We are grateful to C. Girard and S. Ramakrishnan for assistance with irradiation experiments and generation of sequencing libraries, W. Zhang for crosses performed early in the analysis of the *me6ts* mutant, and David Paul for discussions regarding object-based colocalization analysis. We acknowledge N. Bhatla for creating the Exon-Intron Graphic Maker (wormweb.org) used for Figure 1B. We thank A. Dernburg and V. Jantsch for antibodies and the Caenorhabditis Genetics Center (funded by NIH Office of Research Infrastructure Programs P40 OD010440) for strains. This work was supported by an American Cancer Society Research Professor Award (RP-15-209-01-DDC) and NIH grants R01GM53804 and R35GM126964 to AMV, by a Blavatnik Family Foundation Fellowship and a Stanford Mason Case Fellowship to AH, by an FWF Erwin Schrödinger Fellowship (J-3676) to AW, by a Stanford Graduate Fellowship to KY, by grants from the Taiwan Ministry of Science and Technology (107-2923-B-002-001-MY4 and MoST 104-2628-B-002-002-MY3) to PC, and by grant 1S10OD01227601 from the NCRR to the Stanford Cell Sciences Imaging Facility.

---

**Supplemental Table 1.**
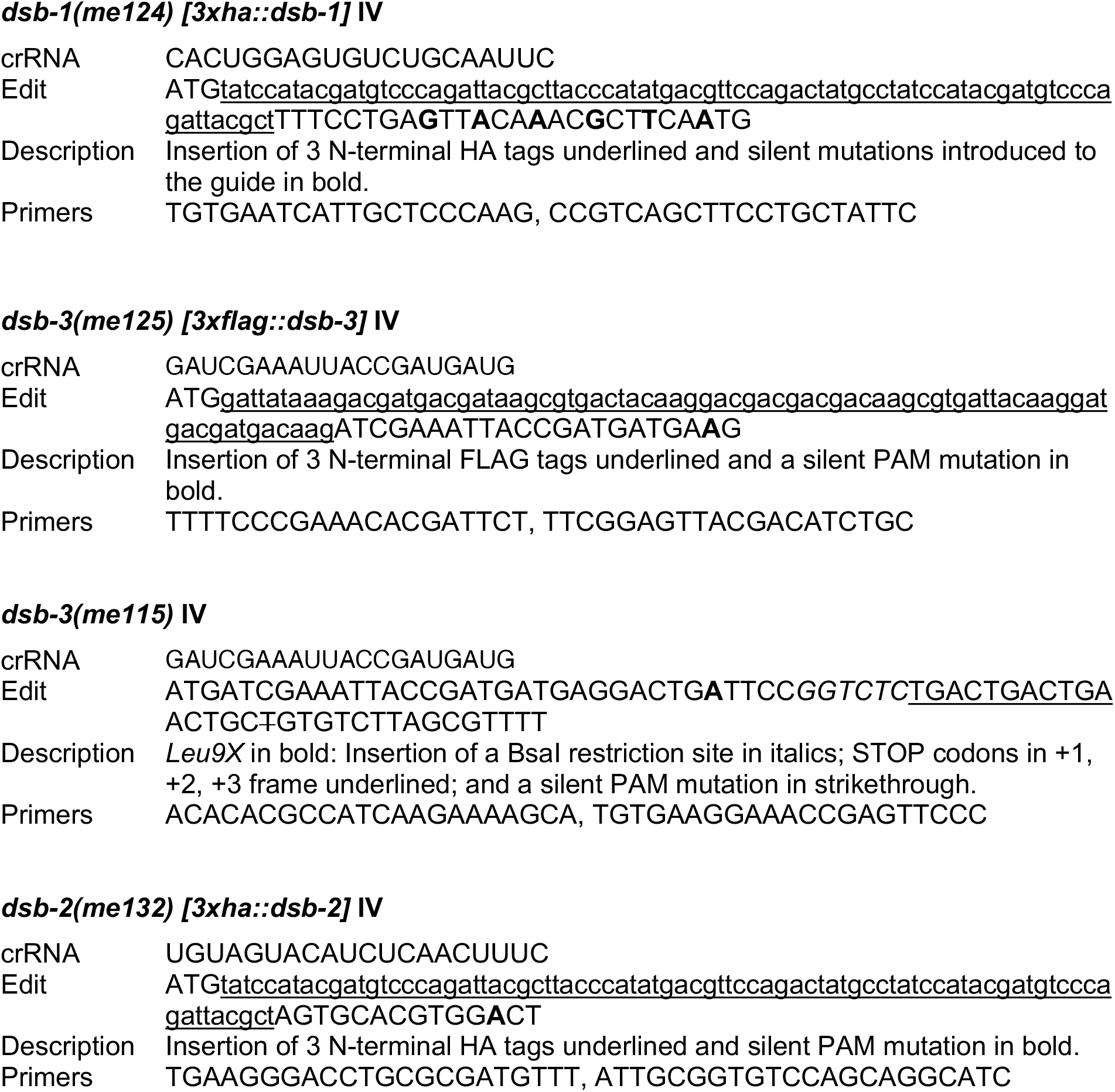
Summary of information relevant to the CRISPR/Cas9 edits created for this work. Information includes: a) sequence of the cRNA used in the injection mixture; b) sequence of the edit created and description of its effect on the gene and encoded protein; c) sequences of primers used for mutation detection and verification of edits via Sanger sequencing.

**Supplemental Figure 1.**
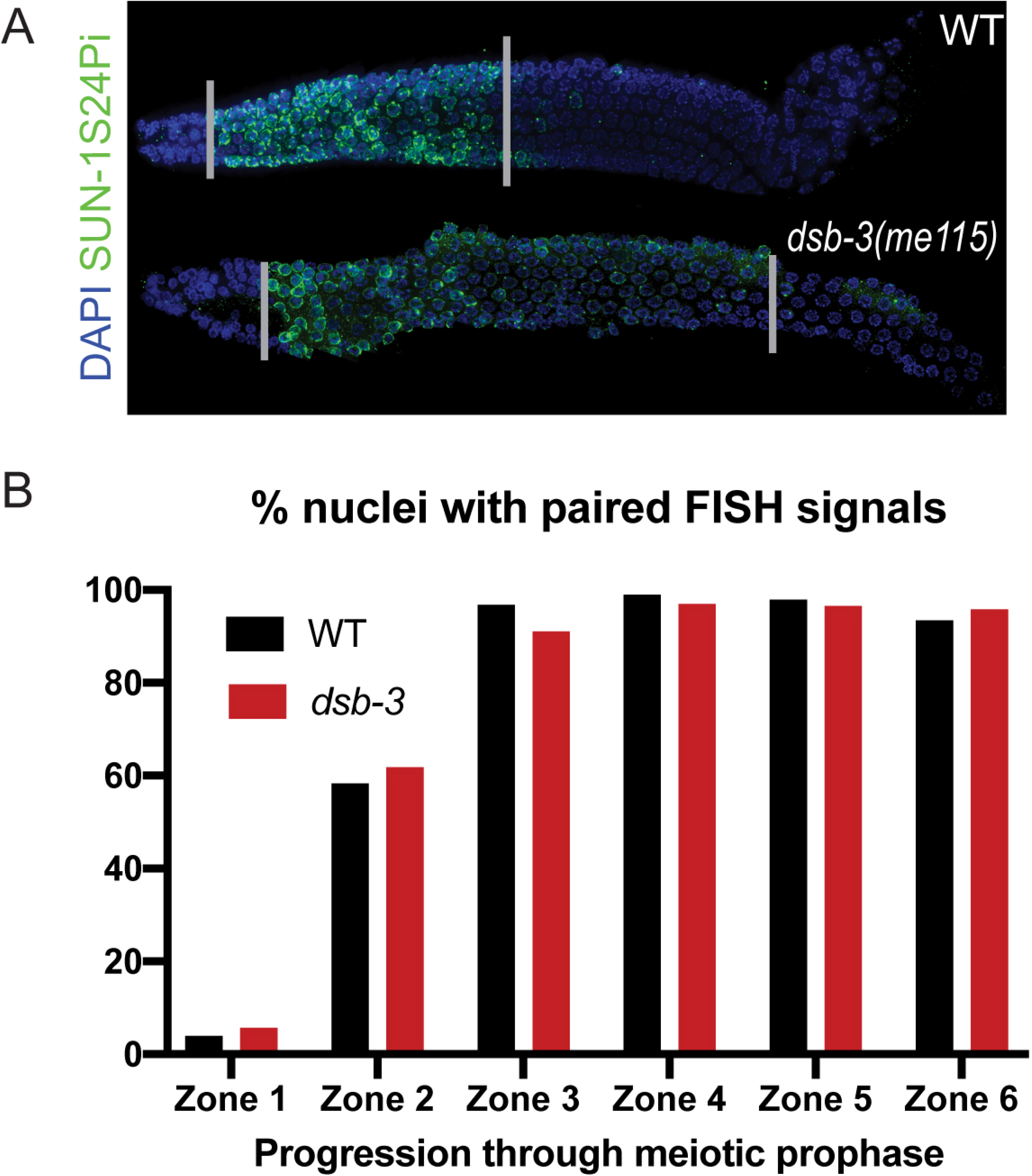
Meiotic prophase progression and homolog pairing in the *dsb-3(null)* mutant. (A) Immunofluorescence images of whole-mount hermaphrodite gonads (from distal tip to end of pachytene) stained with DAPI and antibodies detecting SUN-1 Ser24 Pi, an indicator of CHK-2 activity detected from the onset of meiotic prophase through the early pachytene stage (36, 72). *dsb-3(me115)* mutant germlines shown an extension of this marker relative to WT, reflecting operation of a crossover assurance checkpoint/surveillance mechanism that prolongs the early pachytene stage in response to one or more chromosome pairs lacking crossover-competent recombination intermediates (20, 21, 36). (B) Quantification of homolog pairing assayed by FISH. FISH signals from homologous chromosomes were considered paired if they were separated by ≤ 0.7 μm. Numbers of nuclei scored for WT: zone 1, n=180; zone 2, n=201; zone 3, n=180; zone 4, n=178; zone 5, n=180; zone 6, n=167. Numbers of nuclei scored for *dsb-3*: zone 1, n=159; zone 2, n=180; zone 3, n=155; zone 4, n=160; zone 5, n=143; zone 6, n=117

**Supplemental Figure 2.**
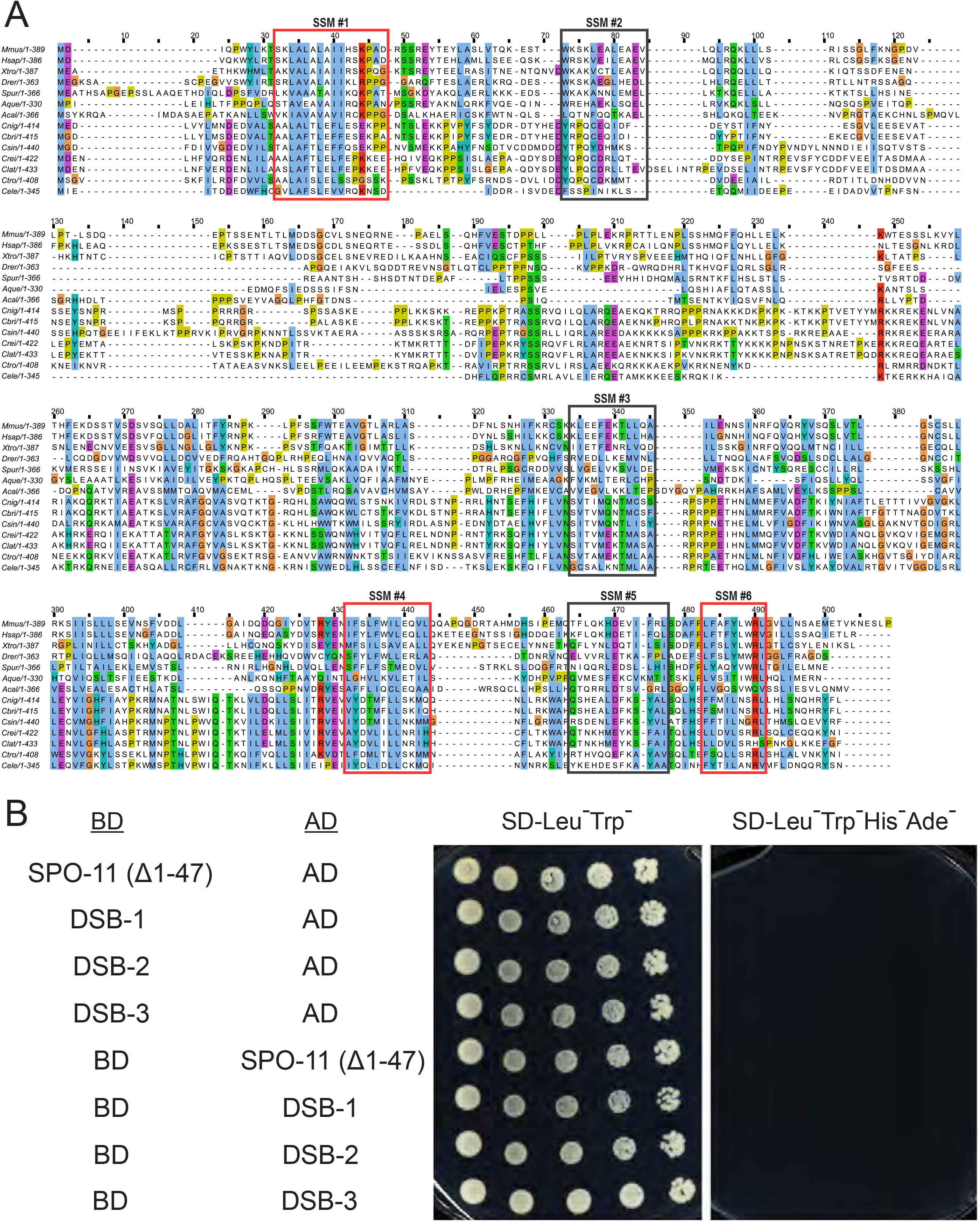
Alignment of MEI4 and DSB-3 orthologs, and Y2H controls. (A) Multiple sequence alignment generated with MAFFT Version 7.0 with the ClustalX coloring scheme. Protein sequences included in the multiple sequence alignment were from the following species: Vertebrates: *Mus musculus, Homo sapiens, Xenopus tropicalis, Danio rerio*; marine invertebrates: *Strongylocentrotus purpuratus, Amphimedon queenslandica, Aplysia californica*; Nematodes of the genus *Caenorhabditis*: *C. nigoni, C. briggsae, C. sinica, C. remanei, C. latens, C. tropicalis, C. elegans*. The outlined boxes indicate the positions of the SSMs that were previously defined for MEI4 orthologs from diverse species. Red boxes indicate cases where the SSM includes at least 5 amino acid residues that are conserved or exhibit similar electrophysiological properties in at least 85% of the aligned sequences. Gray boxes indicate cases where these thresholds are not met. (B) Negative controls for yeast two-hybrid assays, showing lack of auto-activation for cells containing the indicated constructs.

**Supplemental Figure 3.**
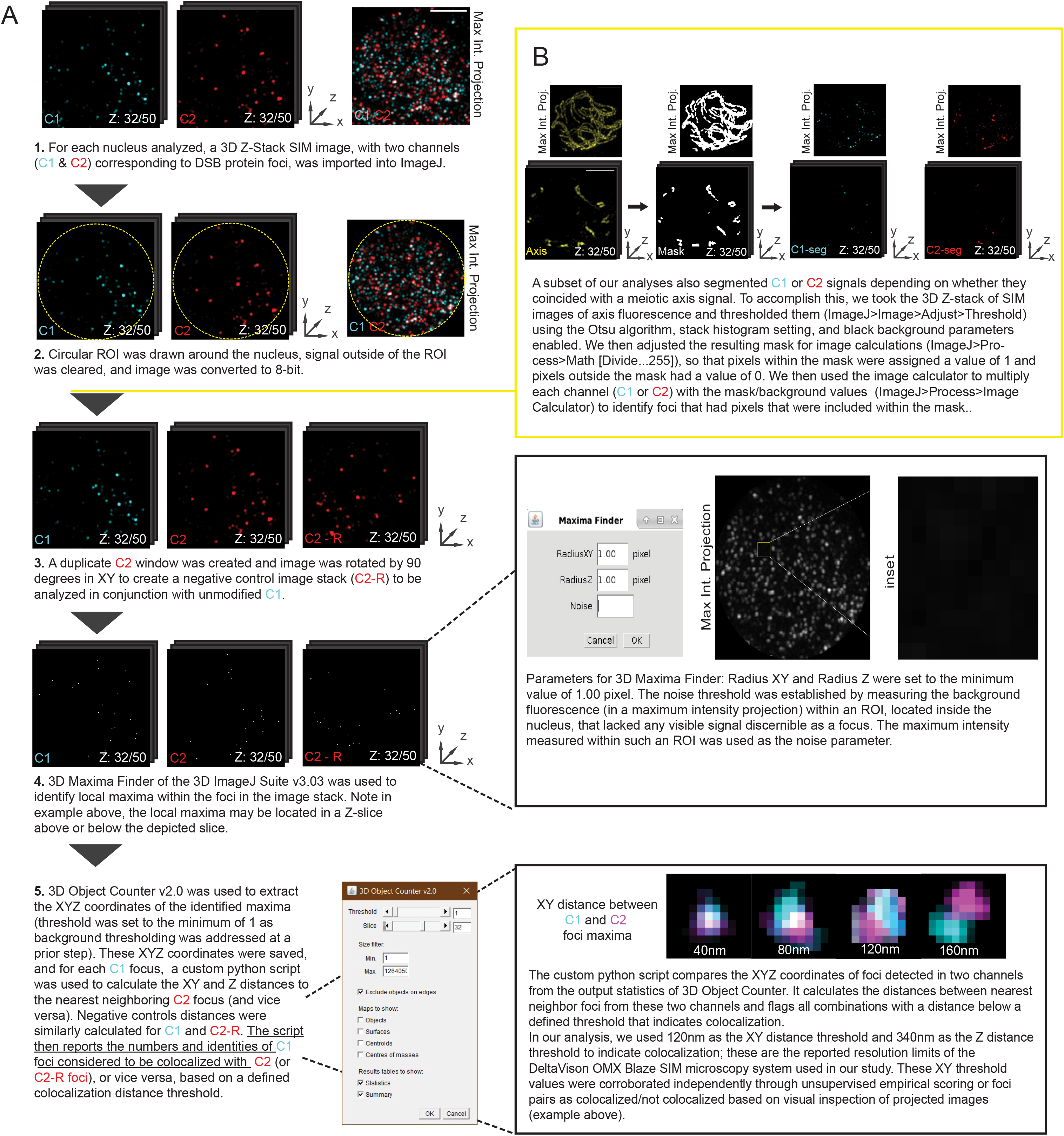
3D Object-Based Colocalization Analysis Pipeline for SIM images of DSB protein foci in spread nuclei. (A) A schematic showing the general pipeline used for colocalization analyses. ImageJ plugins used were 3D Maxima Finder (75) and the 3D Object Counter (81). (B) Yellow box indicating protocol used in a subset of our analyses in which images were additionally segmented to identify DSB protein foci that coincided with the axis signal; the Otsu method (82) was used in our thresholding process for this segmentation.

**Supplemental Figure 4.**
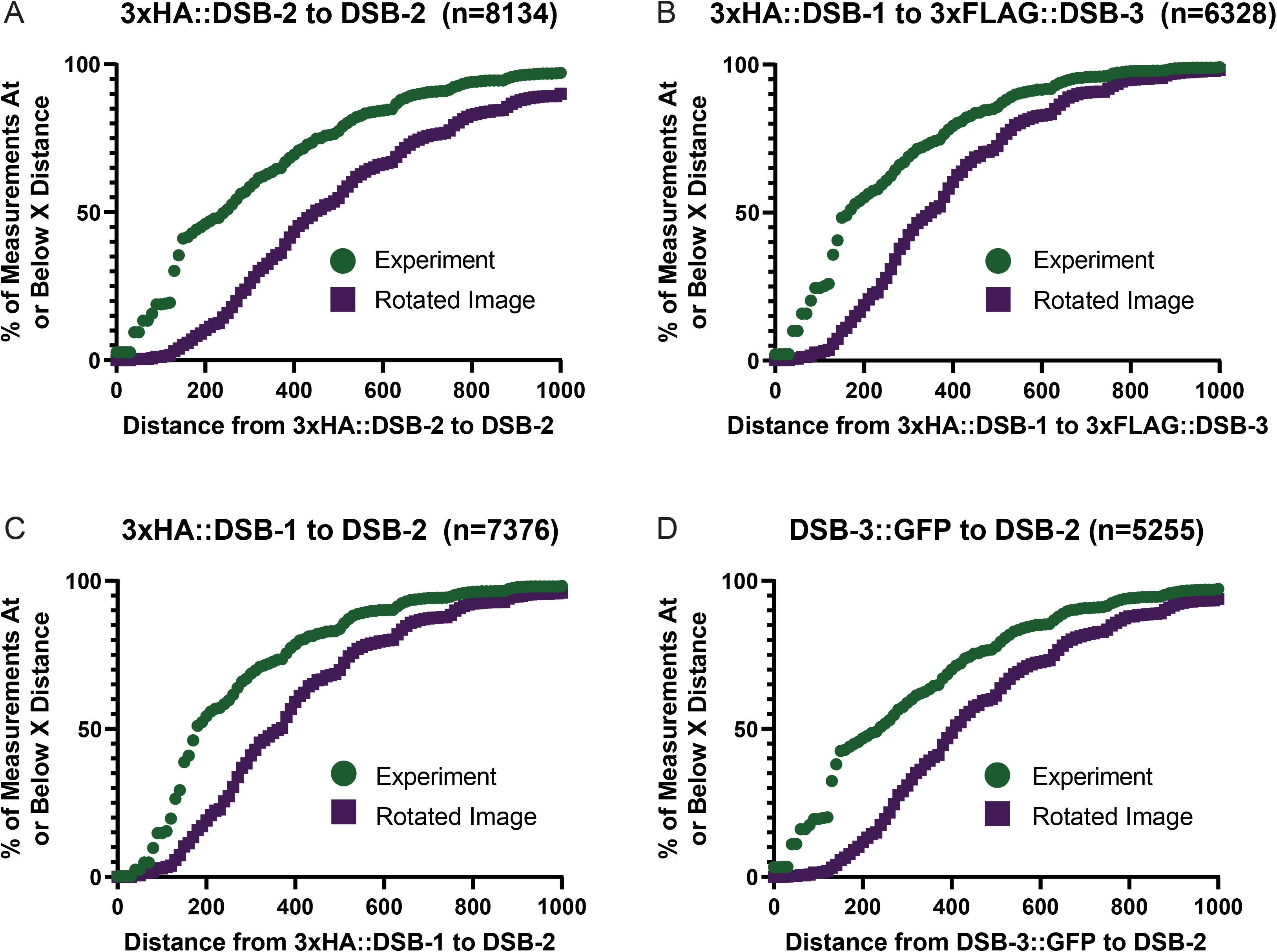
Cumulative distribution plots for the distances from each Channel 1 DSB protein focus to its nearest neighbor Channel 2 focus. The x-axis represents the distances between nearest neighbor foci pairs, and the y-axis indicates the percentage of measurements at or below the given distance on the x-axis. Experimental data are depicted in green circles and values for the corresponding negative control rotated images are indicated with purple squares. (A) The cumulative distribution of distances between 3xHA::DSB-2 mAB foci and nearest neighbor DSB-2 pAB foci. (B) The cumulative distribution of distances between 3xHA::DSB-1 foci and nearest neighbor 3xFLAG::DSB-3 foci. (C) The cumulative distribution of distances between 3xHA::DSB-1 foci and nearest neighbor DSB-2 pAB foci. (D) The cumulative distribution of distances between DSB-3::GFP foci and nearest neighbor DSB-2 pAB foci.

**Supplemental Figure 5.**
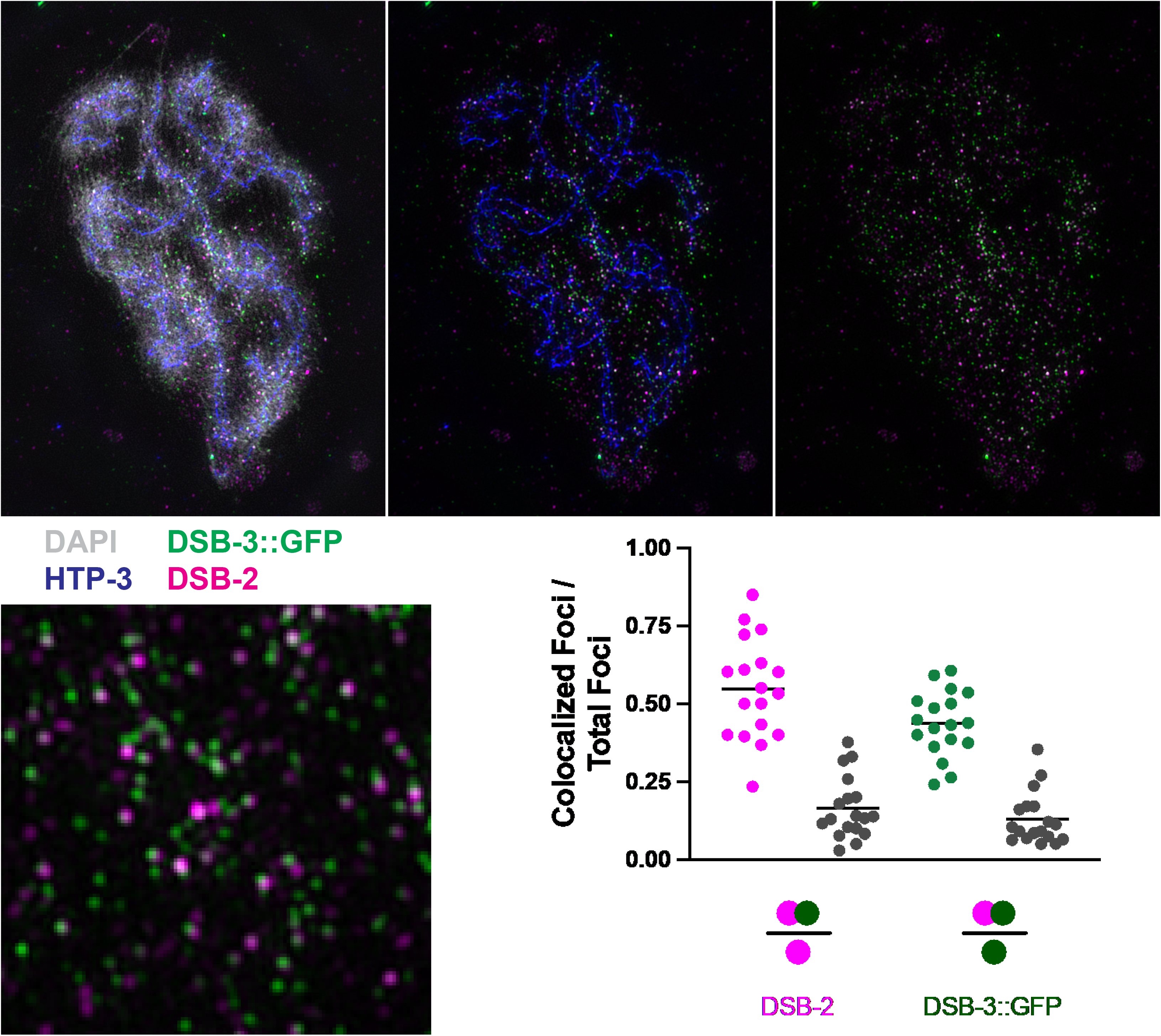
Colocalization analysis for DSB-2 and DSB-3::GFP on super-spread nuclei. Images depict a super-spread nucleus (top) and an inset from the same nucleus (bottom), showing DAPI-stained DNA and immunofluorescence signals corresponding to DSB-2, DSB-3::GFP, and axis protein HTP-3. The graph shows the fraction of DSB-2 foci (magenta) within a given ROI that are colocalized with a DSB-3::GFP focus (green), and vice versa, together with their paired negative controls (represented by grey data points).

